# Live birth protects the developing progeny of the extremophile nematode *Tokorhabditis tufae*

**DOI:** 10.64898/2026.11.15.765984

**Authors:** James Siho Lee, Sara Omer, Cramer Gotthelf, Hyunsoo Yim, Tatsuya Yamashita, Ryoji Shinya, Taisei Kikuchi, Cornelia I. Bargmann

## Abstract

Animals that naturally thrive in extreme environments can yield insights into strategies of resilience to toxic conditions. One such animal is *Tokorhabditis tufae*, a nematode species from Mono Lake, a hypersaline and arsenic-rich lake incompatible with most animal life. Most nematodes reproduce by laying eggs, but *T. tufae* gestates its progeny and gives birth to live young. Here we characterize physiological, anatomical, and behavioral alterations that differentiate *T. tufae* from egg-laying nematodes and contribute to its resilience. We find that progeny are gestated in *T. tufae* through hatching and two larval stages *in utero* before emerging as stress-resistant third-stage dauer larvae. Larvae from the second larval stage are viable after removal from the mother but are killed by high levels of salt and arsenic; gestation of larvae protects them from these stresses. Live birth in *T. tufae* is supported by multiple adaptations to the egg-laying circuit. Canonical serotonergic HSN neurons that drive nematode egg laying are not present in *T. tufae*, and the egg-laying musculature is remodeled into a sphincter-like structure. Live birth is also driven by stage-specific behaviors; *T. tufae* larvae are quiescent during the first two larval stages *in utero* but become active when they reach the dauer stage. The dauer-stage larvae facilitate birth through their own locomotion, linking live birth to physiological stress resistance. These findings reveal developmental and neural adaptations that promote survival in extreme environments.

## Introduction

Environmental contamination by toxic elements is increasing worldwide, posing serious risks to ecological systems and human health (Iqubal et al., 2020; Lay & Levina, 2013; Saaristo et al., 2018). One significant contaminant is arsenic: 230 million humans are exposed to harmful levels of arsenic from sources such as industrial pollution, mining, and agriculture, with symptoms including neuropathy, cancer, and death (Mandal, 2017; Mochizuki, 2019; Prakash et al., 2016; Shaji et al., 2021; Vahter, 2009).

Nematodes are ideal models for studying resilience to environmental stressors, due to their wide distribution and adaptation to harsh environments including arid deserts, the deep sea, mineral-extreme mines, hypersaline lakes, and Siberian permafrost (De Santiago et al., 2026; Jung et al., 2024; Sapir, 2021; Shatilovich et al., 2023; Tintori et al., 2024). We previously described the isolation of the extremophile nematode *Tokorhabditis tufae* from Mono Lake (California, USA), an alkaline lake with high salinity and arsenic levels (Shih, Lee, Shinya, et al., 2019). *T. tufae* adults can survive acute exposure to concentrations of arsenic 150-fold higher than the lethal dose for human cells (Mochizuki, 2019). *T. tufae* and the related nematode species *Auanema freiburgense* and *Auanema rhodense* are also more resistant to arsenite (As(III)) than the laboratory nematode *Caenorhabditis* elegans (Shih, Lee, Shinya, et al., 2019). Their arsenic resistance is supported by variations in the gene encoding DBT-1, a component of branched-chain amino acid catabolism and a target of arsenic toxicity, and is likely to include other factors as well (Shih, Lee, Shinya, et al., 2019; Zdraljevic et al., 2019).

While most nematodes lay their progeny as eggs, *T. tufae* gestates its progeny (Shih, Lee, Shinya, et al., 2019). Here, we show that *T. tufae* protects its progeny from toxic conditions through prolonged gestation across two larval stages before they are born in the third stage as stress-resistant dauer larvae. The second-stage larvae are viable after dissection from the mother, but they are killed by Mono Lake salinity and arsenic. Gestation is facilitated by neuronal and muscular rearrangements of the canonical nematode egg-laying circuit. Live birth is supported by a transition to active locomotion in the emerging dauer larvae, such that birth occurs once progeny are stress-resistant and capable of tolerating the harsh environment of Mono Lake.

## Results

### Prolonged gestation of progeny

Unlike most nematodes, *T. tufae* mothers hatch their eggs internally, gestate larvae in their uterus, and give birth to well-developed larvae (Kanzaki et al., 2021; Yamashita et al., 2023) (**Figure 1A**, **Supplemental Video 1**). At their reproductive peak (day 2 of adulthood), *T. tufae* hermaphrodites hold an average of 26 eggs and 23 larvae *in utero* (**Supplemental Figure 1A**). During live birth, larvae emerge one at a time and always emerge head-first (**Supplemental Figure 1B**). Live births occur at average intervals of 37 minutes (**Supplemental Figure 1C-D**).

**Figure 1.**
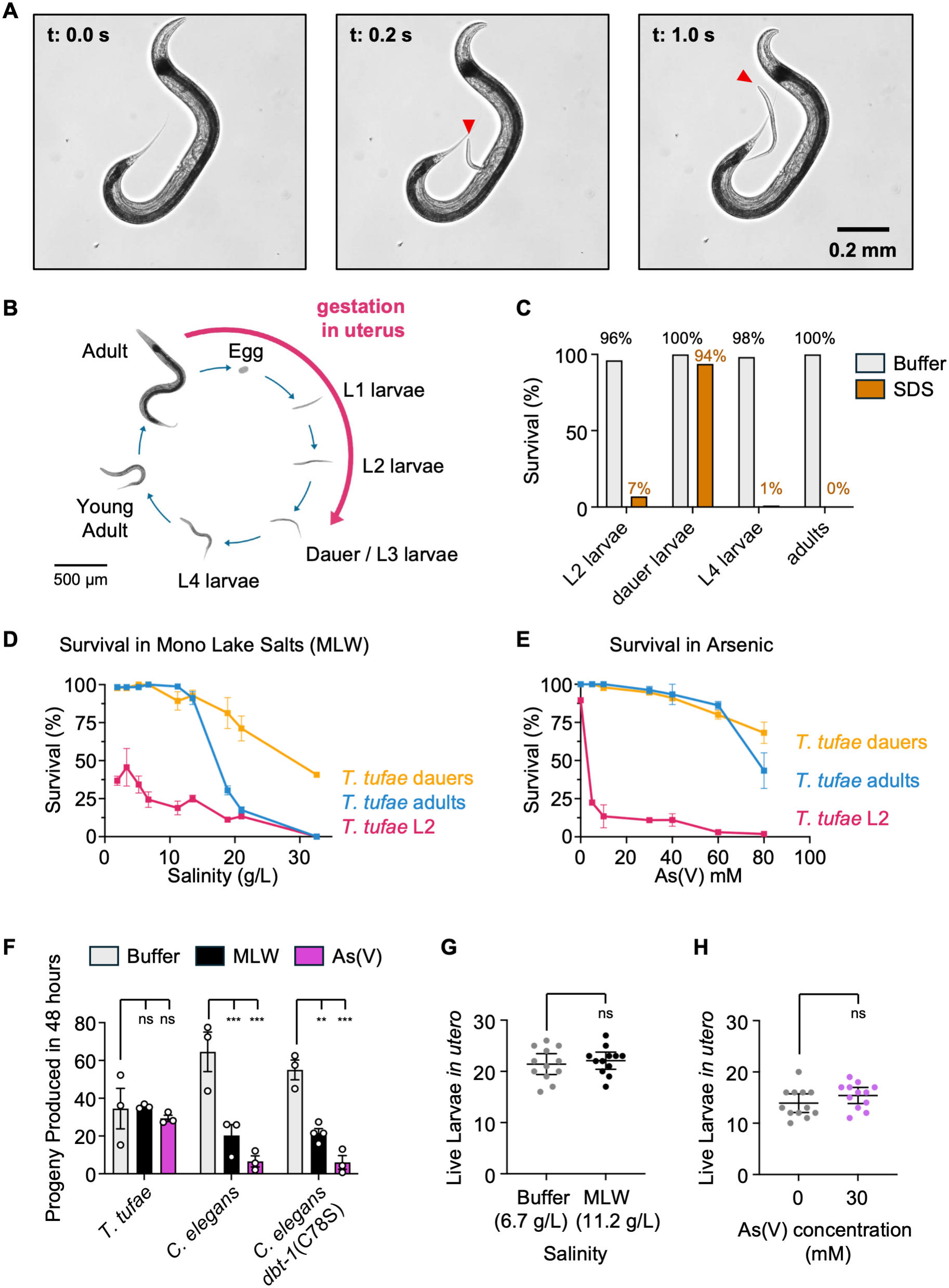
Live birth behavior in *Tokorhabditis tufae* shields progeny from Mono Lake toxins. **(A)** Live birth of a *T. tufae* larva. Red arrowhead indicates the head of the emerging larva. **(B)** Life cycle of *T. tufae*. Gestating progeny progress through the egg, L1, and L2 larval stages *in utero* before being born at the third larval stage (**Supplemental Figures 1** and **2**). **(C)** Survival of dissected *T. tufae* and other stages in NGM buffer (grey) and 1% SDS (orange). n ≥ 60 animals for each condition. **(D-E)** Survival of *T. tufae* after 24-hour exposure to Mono Lake salts (D) or arsenic (E). For each condition, n = 3-6 assays, 10-45 animals per assay. Error bars indicate mean and SEM. **(F)** Progeny produced in 48 hours by *T. tufae*, *C. elegans*, and *C. elegans dbt-1*(C78S) in NGM buffer, 5.2 g/L Mono Lake salinity, and 5 mM arsenic. For each condition, n = 3-6 assays with 10 synchronized adult parents per assay. Error bars indicate mean and SEM. **(G, H)** Survival of *T. tufae* larvae *in utero* after mothers were exposed for 7 hours to (G) NGM buffer or 11.2 g/L Mono Lake salts, and (H) 0 mM or 30 mM arsenic. n = 12 dissected adults. Error bars indicate 95% confidence interval. Statistics: (F) Two-way ANOVA with Tukey’s multiple comparisons test. **p ≤ 0.01; ***p ≤ 0.001; ns, not significantly different. (G, H) unpaired t test.

*T. tufae* progeny are gestated in the uterus through the L1 and L2 larval stages before being born in the third larval stage (**Figure 1B** and **Supplemental Figures 1E-H** and **2**). When gestating progeny are dissected from *T. tufae*, the eggs are inviable, as are small L1 stage larvae that swell upon release. Larger L2 stage larvae are viable and continue development (**Figure 1C** and **Methods**).

**Figure 2.**
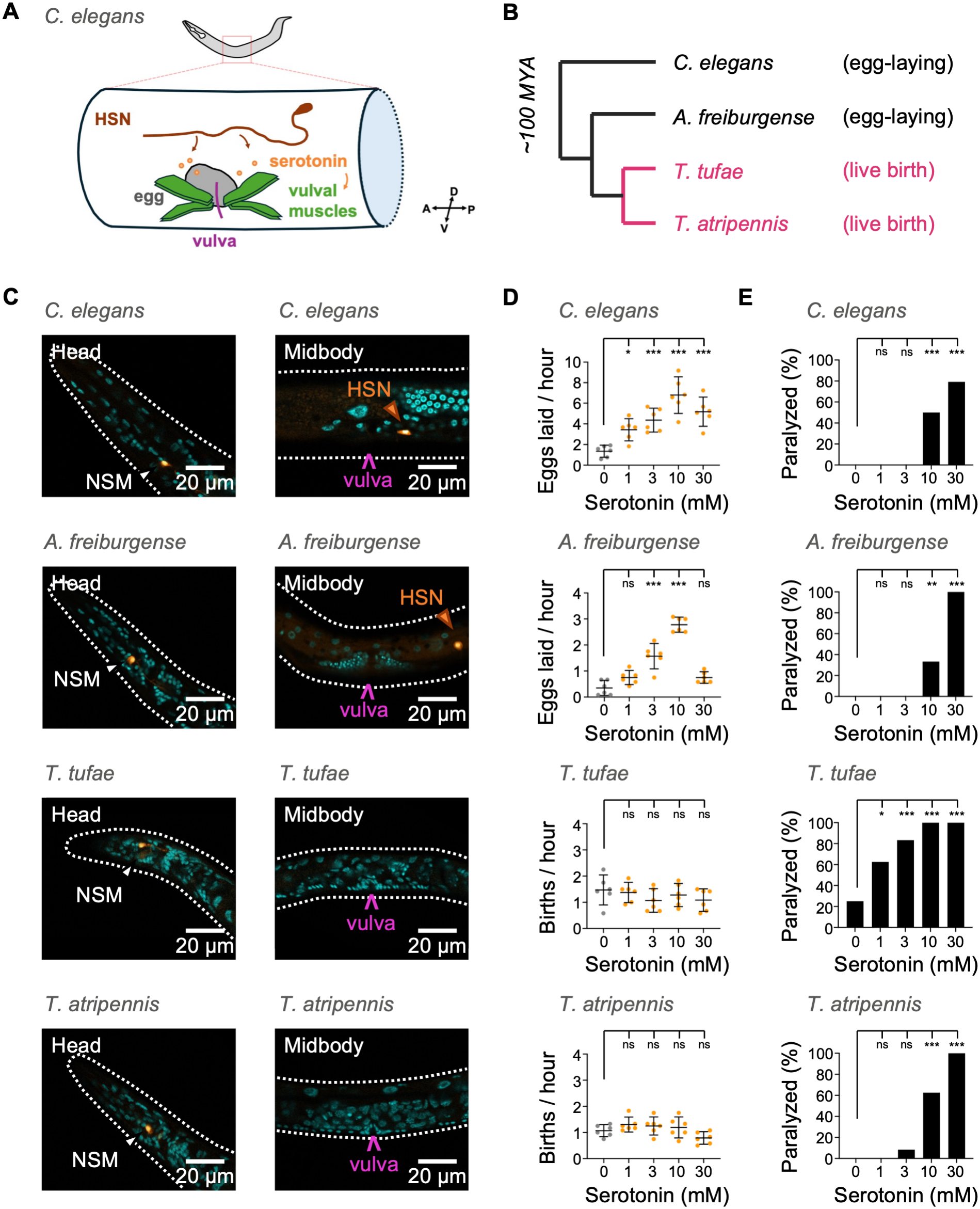
Live birth in *T. tufae* is serotonin-independent. **(A)** Simplified schematic of the *C. elegans* egg-laying circuit. **(B)** Phylogenetic tree showing *Tokorhabditis*, its sister genus *Auanema*, and *Caenorhabditis* and their reproductive modes. **(C)** Serotonin immunostaining indicating the lack of serotonergic HSN in *T. tufae* and *T. atripennis*. (Left) Anti-serotonin (orange) and DAPI (blue) showing presumptive serotonergic NSM feeding neurons (white arrowhead) in the head region of *C. elegans*, *A. freiburgense*, *T. tufae*, and *T. atripennis*. (Right) Serotonergic HSN egg-laying neurons (orange arrowheads) are present in the midbody of *C. elegans* and *A. freiburgense* but not of *T. tufae* and *T. atripennis.* All animals are oriented anterior to the left and ventral down. For each species, images shown are representative of n = 14-30 stained L4s to adults. **(D)** Exogenous serotonin increases egg-laying in *C. elegans* and *A. freiburgense* but does not affect birth rates in *T. tufae* and *T. atripennis*. **(E)** Exogenous serotonin induces locomotor paralysis in all species, indicating bioavailability. For (D-E), n = 6 assays, 4 animals per condition. Error bars indicate 95% confidence interval. Statistics: (D) One-way ANOVA with Dunnett’s multiple comparisons test; (E) Fisher’s exact test. *p ≤ 0.05; **p ≤ 0.01; ***p ≤ 0.001.

The emerging third-stage larvae have two alternate forms that resemble L3 and dauer larvae of *C. elegans,* respectively. *T. tufae* dauers have characteristic dauer morphological features including cuticular rearrangements that seal the mouth, lateral ridges (alae) along the dauer cuticle, and a second cuticular coat over the dauer cuticle (Kanzaki, Kiontke, et al., 2017) (**Supplemental Figure 1I-J**). Like *C. elegans* dauer larvae, which are highly stress-resistant, *T. tufae* dauer larvae are resistant to killing by the harsh detergent sodium dodecyl sulfate (SDS) (Cassada & Russell, 1975; Yamashita et al., 2025) (**Figure 1C**). Dauers undergo two further molts before becoming adult hermaphrodites (**Supplemental Figure 2**). An alternative third larval form in *T. tufae*, the L3 larva, is infrequent and will not be discussed further.

### Prolonged gestation protects progeny from Mono Lake toxins

Because Mono Lake has freshwater inlets but no outlets, the lake has sharp gradients of salts enriched in carbonate, sodium, sulfate, and boron (Jones & Stokes Associates Inc., 1993). *T. tufae* was isolated from a region of the lake that is fed by freshwater streams and ranges from 0.2 to 5.2 g/L in salinity, but adjacent regions of the lake can reach 94.4 g/L salinity (Jones & Stokes Associates Inc., 1993; Shih, Lee, Shinya, et al., 2019). Elevated levels of 200 μM arsenic are present in the lake as well, predominantly as As(V) arsenate at the lake’s surface where nematodes can be found (Oremland et al., 2004; Shih, Lee, Shinya, et al., 2019).

In *C. elegans,* osmotic stress induces egg retention and internal hatching of larvae, which protects the developing progeny from environmental stress (Chen & Caswell-Chen, 2004; Mignerot et al., 2024; Yoon et al., 2020). We hypothesized that prolonged gestation in *T. tufae* may shield developing offspring from the toxic conditions of Mono Lake. To investigate this hypothesis, we compared the sensitivity of adults, dauers, and gestating progeny to Mono Lake salinity and arsenic. Dauers were more resistant to salinity and arsenic compared to other stages, L2 larvae were highly sensitive, and adults had intermediate sensitivity (**Figure 1D-E**). These results indicate that gestating progeny are sensitive to salinity and arsenic but acquire resistance by the time they are born as dauers. Notably, dauers and adults survived in salinity and arsenic levels exceeding those at the site where *T. tufae* was originally isolated.

To further characterize the stress tolerances of *T. tufae*, we compared the survival of *T. tufae* adults, dauers, and L2 to equivalent stages of *C. elegans. C. elegans* can survive across a range of salinity and arsenic levels (Burton et al., 2017, 2021; Khanna et al., 1997; Liao & Yu, 2005). Some natural variants of *C. elegans* have higher arsenic tolerance due to a protective allele in the branched-chain α-keto acid dehydrogenase E2 subunit, *dbt-1*(C78S) (Zdraljevic et al., 2019), which is present in *T. tufae* (Shih, Lee, Shinya, et al., 2019). We found that *T. tufae* adults could tolerate higher levels of salinity and arsenic than *C. elegans* adults, and similar levels of arsenic as *C. elegans* adults with the protective *dbt-1*(C78S) allele (**Supplemental Figure 3A, 3E**). Dauers of both species demonstrated robust tolerance to high salinity and arsenic (**Supplemental Figure 3B, 3F**), whereas the L2 larvae of both species were more sensitive to high salinity and arsenic (**Supplemental Figure 3C, 3G**). These comparisons highlight the resilience of *T. tufae* adults, as well as dauers of both species to Mono Lake toxins.

We hypothesized that gestation might allow *T. tufae* to protect progeny from high salt and arsenic until they emerge as stress-tolerant dauers. To investigate this possibility, we compared population growth of *T. tufae* and *C. elegans* in Mono Lake salinity and arsenic. Animals were grown on Nematode Growth Media (NGM), 5.2 g/L Mono Lake salinity (the highest salinity recorded at the site where *T. tufae* was isolated), or 5 mM arsenic (exceeding the 200 μM concentration in Mono Lake). *T. tufae* produced a similar number of viable progeny under all conditions, whereas *C. elegans* and *C. elegans dbt-1*(C78S) produced fewer viable progeny in Mono Lake salt and arsenic (**Figure 1F**).

To further confirm that gestation shields developing progeny from Mono Lake toxins, we exposed *T. tufae* hermaphrodites for 7 hours to 11.2 g/L Mono Lake salinity or 30 mM arsenic. These conditions killed dissected *T. tufae* L2 (**Supplemental Figure 3I-J**) but were tolerated by *in utero* larvae, indicating that gestation can protect internal progeny from environmental toxins (**Figure 1G-H**).

### Canonical egg laying circuits do not drive live birth

To understand the adaptations in *T. tufae* that allow live birth, we compared neuromuscular structures associated with birth in the egg-laying nematodes *C. elegans* and *Auanema freiburgense*, a member of the sister genus of *Tokorhabditis* (Kanzaki, Kiontke, et al., 2017), and the live-bearing nematodes *T. tufae* and *Tokorhabditis atripennis* (Ragsdale et al., 2022) (**Figure 2A-B**). *C. elegans* egg laying is stimulated by midbody motor neurons called HSNs that release serotonin, which causes vulval muscles to contract and release eggs (Schafer, 2005). We found that anti-serotonin antibodies labelled a pair of neurons in the midbody of *A. freiburgense* that resembled serotonergic HSNs in *C. elegans*. By contrast, serotonin immunolabelled neurons were absent from the midbody of *T. tufae* and *T. atripennis* (**Figure 2C**, right). All species had serotonin immunolabelleled neurons in two pharyngeal neurons in the head, the presumptive NSM feeding neurons (**Figure 2C**, left) (Horvitz et al., 1982; Rhoades et al., 2019). Therefore, although serotonin is present in all species, it appears to be absent in HSN egg-laying neurons in *Tokorhabditis*.

Exogenous serotonin stimulated egg laying in *C. elegans* and *A. freiburgense* but did not stimulate the emergence of eggs or larvae from *T. tufae* and *T. atripennis* **(Figure 2D**). Exogenous serotonin induced locomotor quiescence in *C. elegans*, *A. freiburgense*, *T. tufae,* and *T. atripennis*, indicating that serotonin was absorbed and biologically active in all species (**Figure 2E**) (Gürel et al., 2012; Ishita et al., 2021). These results suggest that serotonin is both anatomically and functionally uncoupled from birth in the two *Tokorhabditis* species.

In addition to serotonin, egg laying in *C. elegans* is stimulated by acetylcholine secreted from VC motor neurons (Collins et al., 2016; Schafer, 2005) and inhibited by tyramine secreted from neuroendocrine uv1 cells (Alkema et al., 2005; Collins et al., 2016). Live birth in *T. tufae* was unaffected by the cholinergic agonist levamisole or by exogenous tyramine (**Supplemental Figure 4**) (Trent et al., 1983; Weinshenker et al., 1995).

### *T. tufae* lacks canonical HSN egg-laying neurons

Many nematode species have similar body plans, including comparable numbers of neurons and their types, allowing homologous neurons to be identified across species (Han et al., 2016; Walrond et al., 1985). However, certain neurons can be gained, lost, or modified across species lineages (Cook et al., 2024; Loer et al., 2026; Loer & Rivard, 2007; Toker et al., 2025). The lack of anti-serotonin labeling of HSN egg-laying neurons in *T. tufae* is consistent with two general possibilities: (1) HSN neurons are not serotonergic in *T. tufae* or (2) HSN neurons are absent in *T. tufae*.

To investigate these possibilities, we examined the expression of known HSN marker genes in *T. tufae* using HCR-FISH. Tested genes included the *T. tufae* homologs of *tph-1*, which encodes the serotonin biosynthetic enzyme tryptophan hydroxylase (Sze et al., 2000); *unc-86*, which encodes a homeobox transcription factor crucial for the specification of the HSN egg-laying neurons (Hobert, 2021; Sze et al., 2002); and *egl-3*, which encodes a proprotein convertase involved in neuropeptide maturation (Li & Kim, 2008) (**Methods**). These genes are all expressed in the HSN egg-laying and NSM feeding neurons of *C. elegans* (Hobert, 2021; Lloret-Fernández et al., 2018; Taylor et al., 2021). In *T. tufae*, we observed co-expression of *tph-1*, *unc-86*, and *egl-3* in the NSM feeding neurons (**Figure 3A**), but not elsewhere in the body (**Figure 3B**), consistent with the loss of characteristic HSN features.

**Figure 3.**
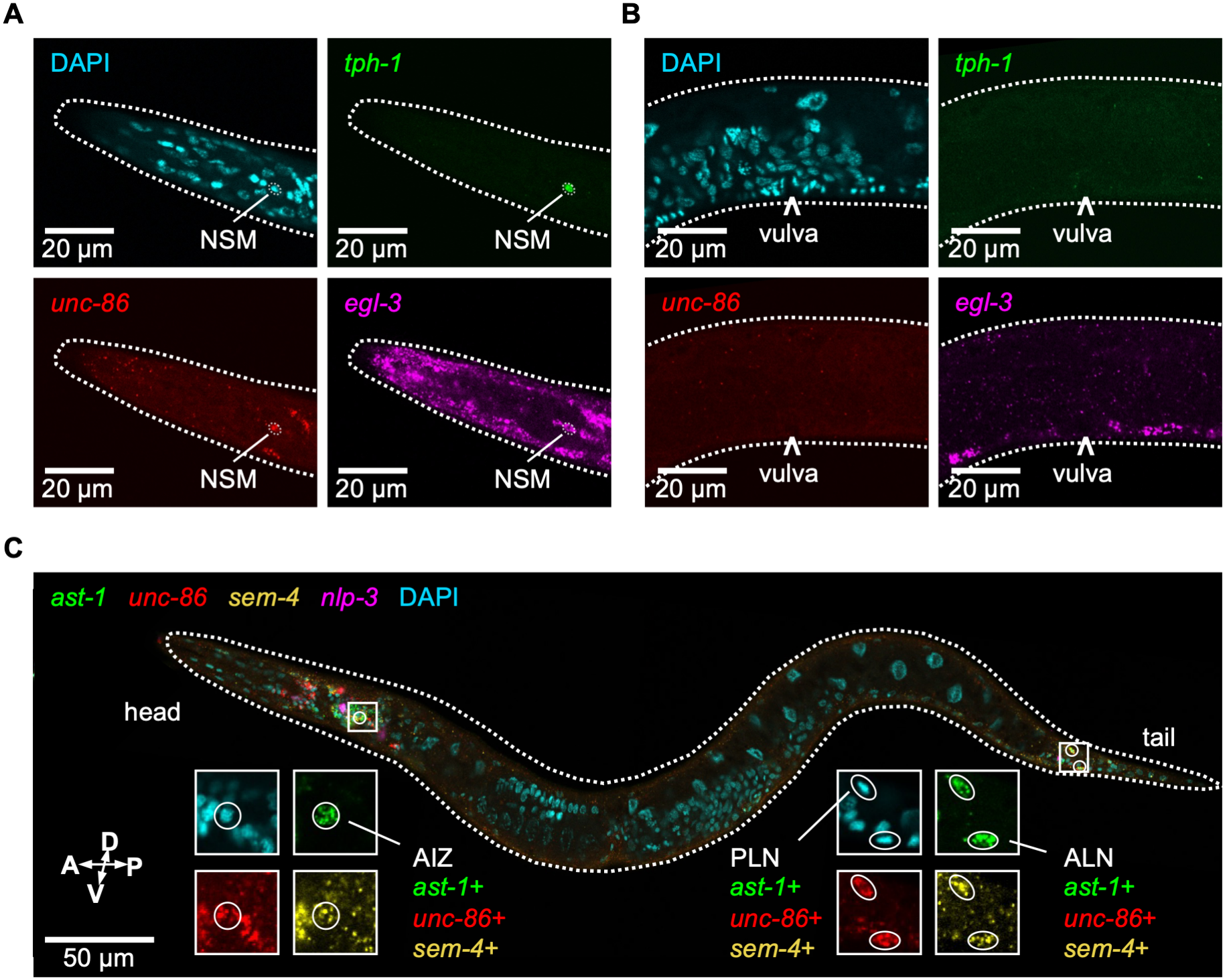
*T. tufae* lacks canonical HSN egg-laying neurons. **(A-B)** Hybridization Chain Reaction Fluorescent *In Situ* Hybridization (HCR-FISH) in *T. tufae* for *tph-1*, *unc-86*, and *egl-3*, markers of NSM and HSN in other nematodes. These genes are co-expressed in the NSM head neuron (A) but not in the midbody (B) or elsewhere (not shown). **(C)** HCR-FISH in *T. tufae* for *ast-1*, *unc-86*, *sem-4*, and *nlp-3,* which are co-expressed in the HSN in *C. elegans.* Co-expression of all four genes is not observed in *T. tufae*. Co-expression of *ast-1*, *unc-86*, and *sem-4* is observed in neurons that may correspond to AIZ head neurons and PLN and ALN tail neurons. All animals are oriented anterior to the left and ventral down. Images shown are representative of 11 stained L4 animals.

We next investigated *ast-1*, *unc-86*, and *sem-4*, which encode transcription factors that coordinately define *C. elegans* HSN identity (Lloret-Fernández et al., 2018), and *nlp-3,* which encodes a neuropeptide secreted by the *C. elegans* HSN (Brewer et al., 2019). These four genes are co-expressed in the *C. elegans* HSN, but they were not coordinately co-expressed in *T. tufae* (**Figure 3C**). Co-expression of *ast-1*, *unc-86*, and *sem-4* is also observed in the *C. elegans* AIZ head neurons and ALN and PLN tail neurons (Olson et al., manuscript in preparation; Taylor et al., 2021, 2026); co-expression of these genes was observed in *T. tufae* in a pair of head neurons (the presumptive AIZ) and two pairs of tail neurons (the presumptive ALN and PLN neurons). Thus, multiple neuronal cell types are shared by *C. elegans* and *T. tufae*, but neurons with an HSN-like molecular profile appear to be absent in *T. tufae*.

### *T. tufae* has distinct muscular specializations for live birth

In *C. elegans*, eggs are released when vulval muscles that connect the vulva to the body wall contract to open the vulva (Collins et al., 2016; Schafer, 2005) (**Figure 4A**). We used actin staining to compare the muscles of the egg-laying nematodes *C. elegans* and *A. freiburgense*, and the live-bearing nematodes *T. tufae* and *T. atripennis*. *C. elegans* vulval muscles have a characteristic X-shaped arrangement at the vulva (**Figure 4A-B**); a similar geometry is present in *A. freiburgense* and in another *Auanema* species, *A. rhodense* (**Figure 4B** and **Supplemental Figure 5**). By contrast, vulval muscles in *T. tufae* and *T. atripennis* formed an O-shaped radial ring around the vulval opening (**Figure 4A-B**).

**Figure 4.**
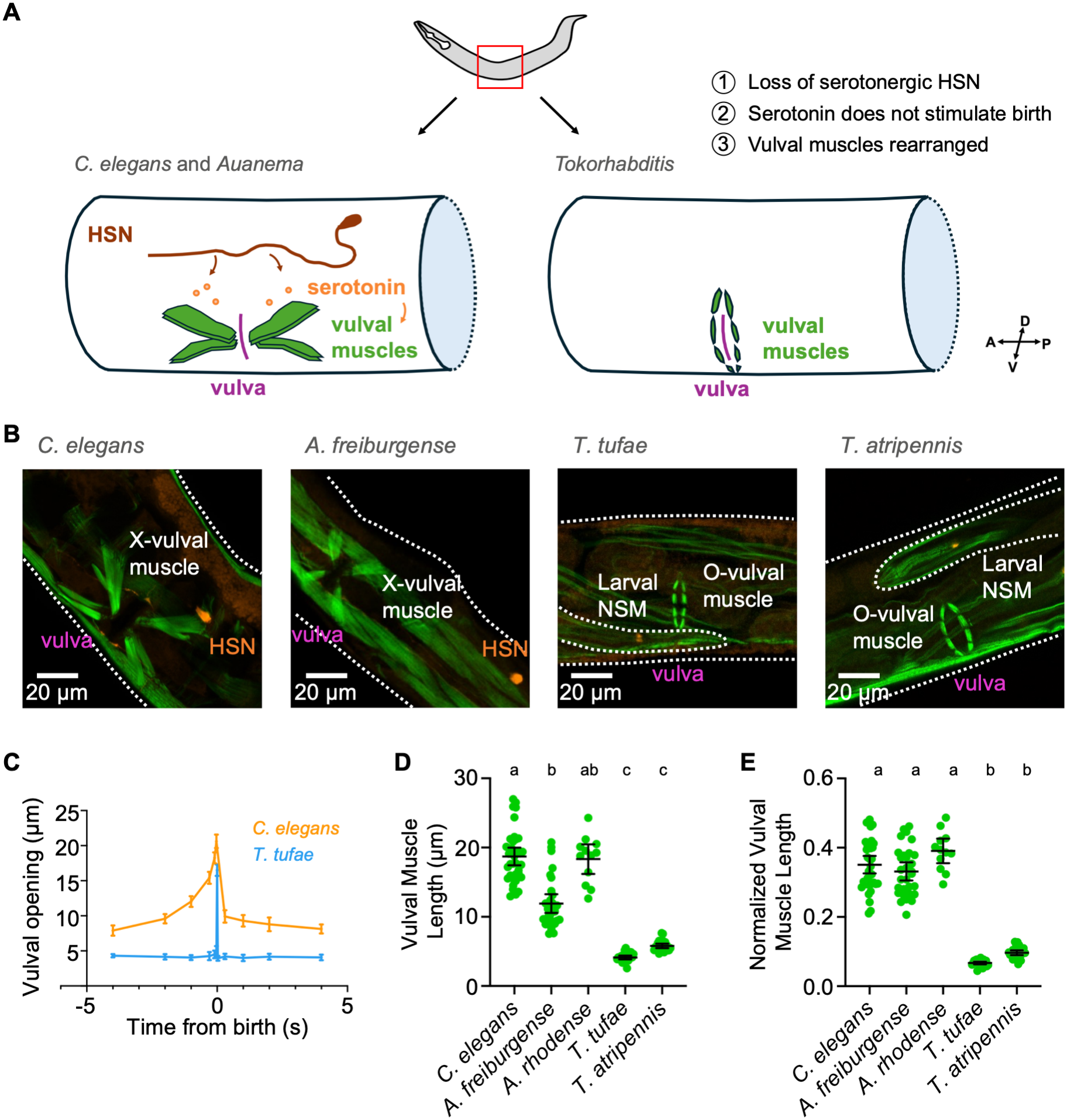
Vulval muscle specializations across species. **(A)** Schematic comparing vulval structures of *C. elegans*, *Auanema,* and *Tokorhabditis*. **(B)** Antibody staining showing muscles (actin, green) and serotonin (orange) in the midbody of *C. elegans*, *A. freiburgense*, *T. tufae*, and *T. atripennis*. Animals are oriented with anterior at left; ventral views highlight X-shaped vulval muscles in *C. elegans* and *A. freiburgense* and O-shaped vulval muscles in *T. tufae* and *T. atripennis*. For clarity, a single *in utero* larva is outlined in *T. tufae* and *T. atripennis*; in each case, its NSM neuron (orange) is serotonin-positive. Images shown are representative of n = 14-26 animals per species. **(C)** Opening of the vulva, measured as the distance between the vulval lips, precedes *C. elegans* egg-laying but does not precede *T. tufae* live birth. For *C. elegans*, n = 31 events from 16 mothers; *T. tufae*, n = 20 events from 15 mothers. Error bars indicate 95% confidence interval. **(D-E)** Length of individual vulval muscle cells in *C. elegans*, *A. freiburgense*, *A. rhodense*, *T. tufae*, and *T. atripennis*. (D) Raw lengths and (E) lengths normalized to body width. n = 12-36 vulval muscles measured from 6-8 adults. Error bars indicate 95% confidence interval. Statistics: (D-E) Kruskal-Wallis test with Dunn’s multiple comparisons test. Letters indicate *q*-value groups.

During egg-laying, *C. elegans* vulval muscle contraction opens the vulva over a few seconds to allow the release of eggs (Collins et al., 2016) (**Figure 4C**). Vulval opening in *T. tufae* was limited to < 0.1 seconds at the moment of larval birth (**Figure 4C** and **Supplemental Video 2**). Together with the smaller size of the vulval muscles in *Tokorhabditis* at rest (**Figure 4D-E**), these dynamics suggest that the muscle ring of *T. tufae* acts as a sphincter that relaxes at birth.

### Larval behaviors support live birth

High-resolution video recordings demonstrated that *T. tufae* larvae were behaviorally quiescent during development in the uterus, with a straight posture reminiscent of sleep states in *C. elegans* (Hill et al., 2014; Raizen et al., 2008; Tramm et al., 2014) (**Figure 5A**). Immediately before birth, larvae assumed an S-shaped posture and moved in the uterus with their heads positioned at the mother’s vulva (**Figure 5B-C**, **Supplemental Video 3**). The viable L2 larvae dissected from the mother recapitulated these *in utero* dynamics, transitioning suddenly from quiescence to an active state at intervals of 110 minutes on average (**Figure 5D** and **Supplemental Figure 6A**) (**Supplemental Video 4**). Transitions to motility occurred until 14 hours after dissection, after which all dissected larvae were motile (**Supplemental Figure 6B**). By staging *ex utero* larvae, we found that viable quiescent larvae were approximately 68% L2 larvae, 18% molting L2 larvae, and 14% dauers, whereas active larvae were all dauers (**Figure 5E**). In agreement with these observations, over 90% of quiescent larvae were killed by exposure to SDS, whereas over 90% of active larvae survived SDS treatment (**Figure 5F**). These results suggest that activity *in utero* is mainly restricted to the dauer stage.

**Figure 5.**
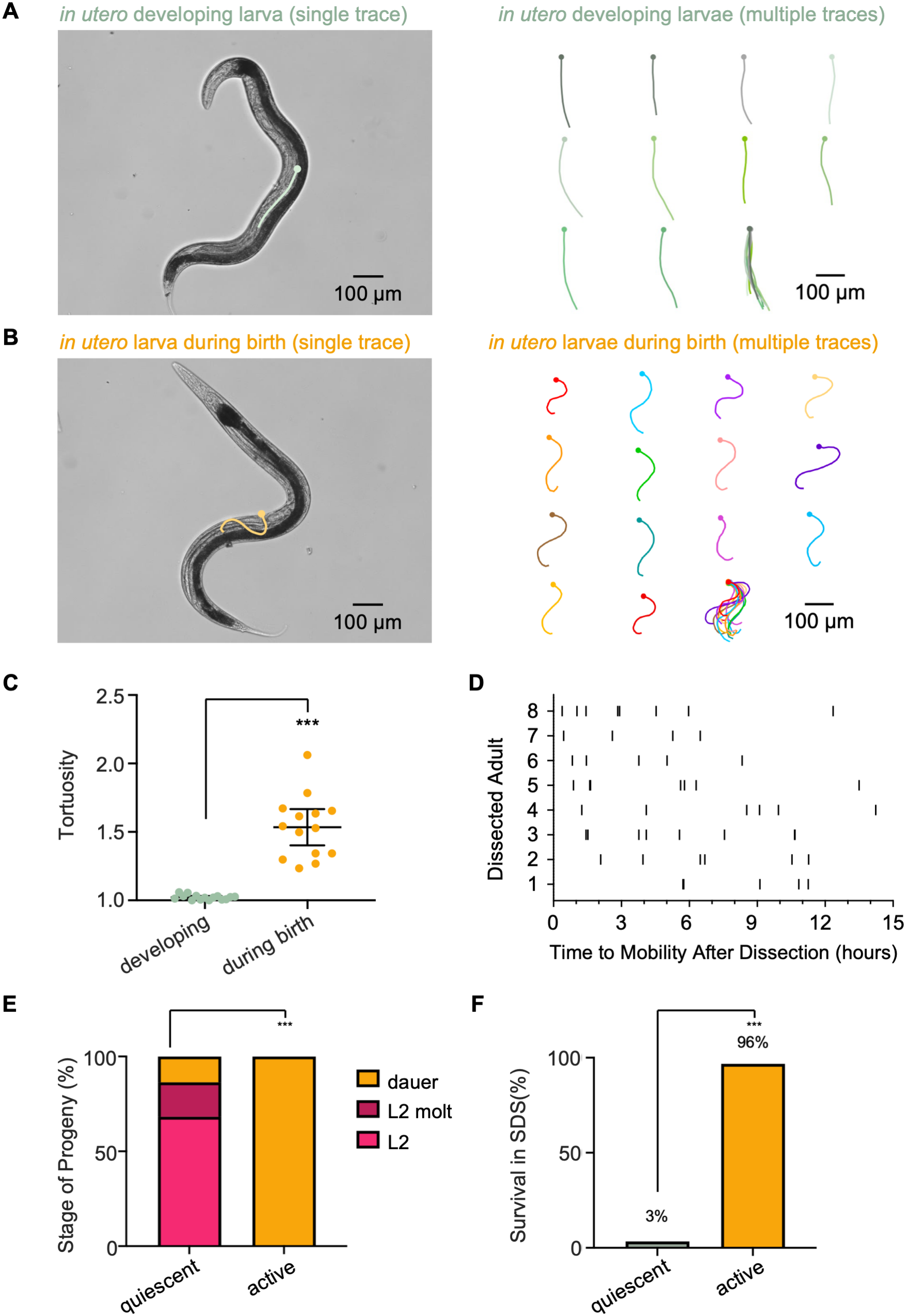
Behavioral specializations in *T. tufae* larvae. **(A)** Early-stage *T. tufae* larvae are quiescent and adopt a straight posture in the uterus. A single larva is traced on the left; traces of 15 larvae from 12 adults are shown on the right. **(B)** Larvae enter a mobile behavioral state during birth. A single larva is traced on the left; traces of 14 larvae from 11 adults are shown on the right. For (A-B), circles indicate the position of the head, and a larval overlay is shown on the lower right. **(C)** Tortuosity measurements of larva postures during development in the uterus or during birth. **(D)** Dissected larvae recapitulate the transition from quiescence to activity *ex utero*. Shown is an ethogram where ticks indicate events in which dissected larvae transition from quiescence to activity. Each tick represents an individual larva dissected from independent adults (rows). n = 49 dissected larvae from 8 adults. **(E)** Developmental stages of dissected progeny (n = 22 quiescent and 21 active larvae). **(F)** SDS-sensitivity of quiescent (n = 59) or active (n = 31) larvae dissected from the uterus.

### *T. tufae* Unc mutants demonstrate impaired live birth

Further insight into live birth emerged from analysis of an uncoordinated (Unc) *T. tufae* mutant isolated after ethyl methanesulfonate (EMS) forward mutagenesis of *T. tufae* (**Supplementary Figure 7**). Locomotion in the Unc mutant was sluggish, uncoordinated, and kinked compared to wild-type animals (**Supplemental Video 5**). The Unc mutant also displayed a 1.9-fold reduction in live birth rate compared to wild type (**Figure 6A**). This reduced birth rate was not due to differences in the number of larvae gestated *in utero*, which was similar between Unc and wild-type animals (**Figure 6B**). Unc mutants died by day 5 of adulthood, a time at which 100% of wild-type adult animals were alive; their death was associated with *in utero* larvae that failed to emerge, a known cause of maternal death in *C. elegans* (**Figure 6C** and **Methods**).

**Figure 6.**
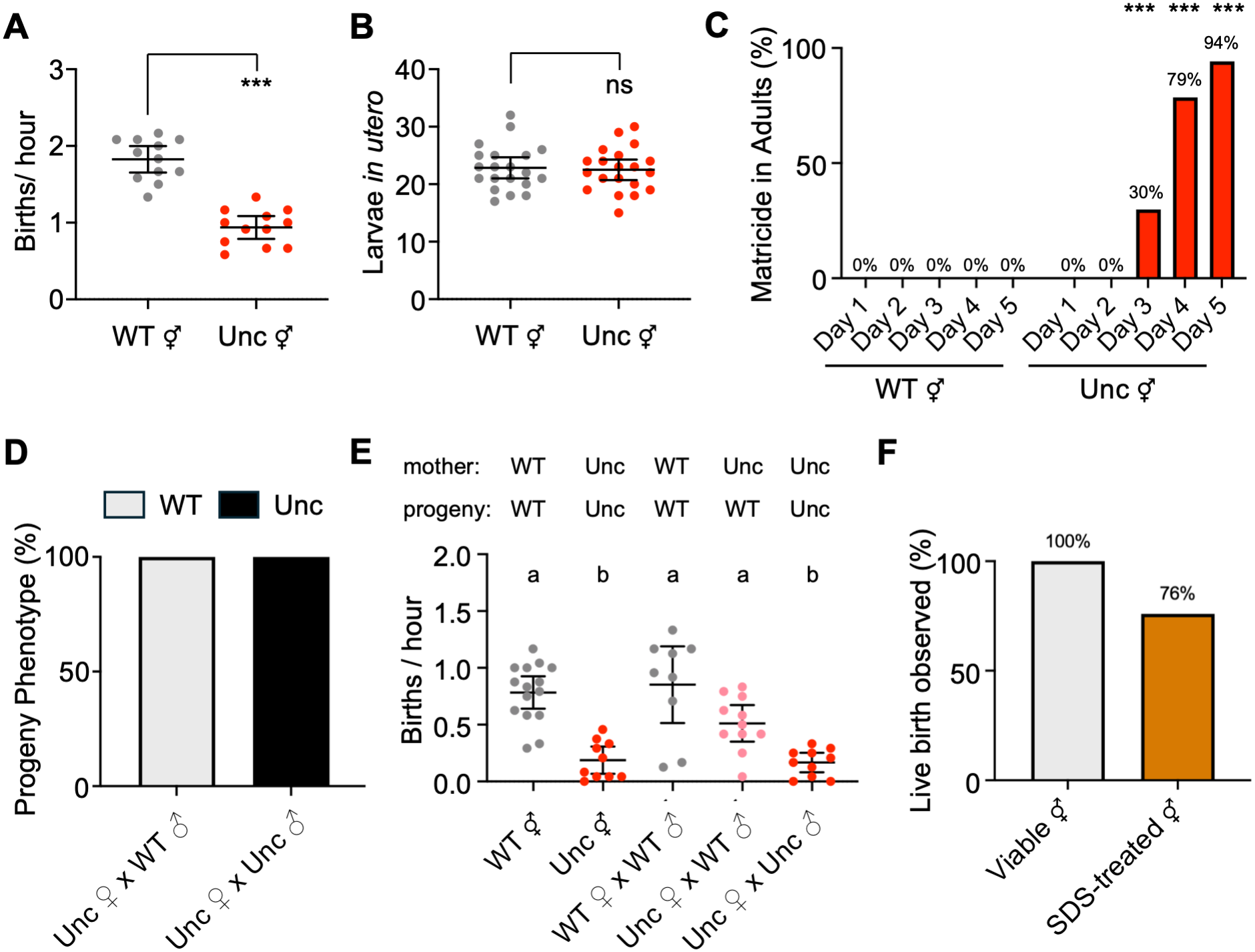
*T. tufae* Unc mutants demonstrate impaired live birth. **(A)** Birth rates in wild type and Unc *T. tufae* (n = 12 assays, 4 animals per assay). **(B)** Number of larvae *in utero* in wild type and Unc mutants (n = 20 adults dissected each). **(C)** Death of adult hermaphrodites, possibly associated with retention of active progeny (n = 80 wild-type adults and 224 Unc adults). **(D)** The Unc phenotype is recessive. n = 134 cross progeny from Unc ♀ x WT ♂ cross; 30 cross progeny from Unc ♀ x Unc ♂ cross. **(E)** Birth rate of wild-type or Unc ♀ mated to wild-type or Unc ♂, and age-matched day-5 adult ⚥. Note that all birth rates are lower in older mothers. Phenotypes of mothers and progeny are indicated (n = 9-15 day-5 adults). **(F)** Percentage of viable or SDS-treated adults observed to give live birth. n = 15 viable adults, 21 SDS-treated adults. Statistics: (A-B) unpaired t test. ***p ≤ 0.001; ns, not significantly different; (C) Fisher’s exact test. ***p ≤ 0.001 for comparisons between wild type and Unc mutants on matched days; (E) Brown-Forsythe ANOVA test with Dunnett’s T3 multiple comparisons test. Letters indicate q-value groups.

To ask whether the impaired birth rate of Unc mutants was caused by defects in the adults or in the gestating larvae, we took advantage of the rare female and male *T. tufae* that arise in each generation (Kanzaki et al., 2021). We first established that when Unc females were mated with wild-type males, the progeny were wild-type, indicating that the Unc is recessive (Unc ♀ x WT ♂ cross) (**Figure 6D**). Next, we determined that, as was true for hermaphrodite birth, the birth rate of Unc progeny from female mothers (Unc ♀ x Unc ♂ cross) was reduced compared to that of wild-type progeny from female mothers (WT ♀ x WT ♂ cross) (**Figure 6E**). Finally, we compared the birth rate of wild-type and Unc progeny that were both born from Unc mothers. Wild-type progeny (Unc ♀ x WT ♂ cross) had higher birth rates than the Unc progeny (Unc ♀ x Unc ♂) (**Figure 6E**). These results suggest that impaired movement of the gestating progeny affects *T. tufae* birth rate.

To better understand the contribution of the mother to larval birth, we investigated whether live birth can occur in *T. tufae* adults killed by treatment with SDS. Brief exposure to SDS killed *T. tufae* adults but spared *in utero* progeny (**Methods**). Live birth was observed in 76% of adults that were killed with SDS (**Figure 6F** and **Supplemental Video 6**), although at a slower rate than in intact animals (**Supplemental Figure 8**). Together, these experiments suggest that both the mother and the larvae contribute to live birth.

## Discussion

Anthropogenic change is driving the emergence of environments enriched in toxic chemicals (Iqubal et al., 2020; Lay & Levina, 2013). Here, we find that *T. tufae* survives toxins associated with its natural environment, Mono Lake, by protecting its developing offspring through live birth behavior. This strategy allows the mothers to shield their young until they develop into stress-resistant dauer larvae. *T. tufae* dauers can withstand Mono Lake levels of arsenic and salt, while developing progeny cannot.

In *T. tufae*, eggs and two stages of juvenile larvae are retained in the mother, and birth is coordinated with dauer development. The dauers are resilient to toxic conditions, are highly active, and develop into self-fertile hermaphrodites that can establish new populations (Bubrig & Fierst, 2021; Yamashita et al., 2023). Thus, dauer offspring are likely well-equipped to survive and disperse in the chemical gradients of Mono Lake (Oremland et al., 2004).

### Adaptations for live birth in nematodes

The phylum of Nematoda contains over 28,000 described species, which nearly all reproduce by egg laying (Hodda, 2022; Kanzaki et al., 2021; Sudhaus & Fitch, 2001). The majority of egg-laying nematode species can retain their eggs for different durations according to environmental conditions, genotype, or species (Mignerot et al., 2024; Sudhaus & Fitch, 2001); some nematode species have eggs that can hatch internally, allowing birth of larvae in simple or facultative forms of live birth (Chen & Caswell-Chen, 2004; Mignerot et al., 2024; Sudhaus & Fitch, 2001). Rare nematode species with advanced live birth demonstrate apparent nutrient provisioning to embryos *in utero,* as inferred by the growth of eggs before they hatch into larvae. Advanced live birth has been directly described in *T. tufae*, whose embryos are permeable to high-molecular-weight substances, a property that could allow the uptake of nutrients *in utero* (Yamashita et al., 2023). Advanced live birth has also been suggested in two other known *Tokorhabditis* species and species in the distantly related genus *Sudhausia* (three species), all of whose embryos increase in size during *in utero* development (Kanzaki et al., 2021; Ragsdale et al., 2022; Herrmann et al., 2013; Kanzaki, Giblin-Davis, et al., 2017).

This spectrum of egg laying to simple and advanced live birth suggests that live birth in nematodes may be supported by several adaptations. Since serotonin expression in the HSN neuron was observed in species of the sister genus *Auanema*, we infer that the loss of serotonergic HSN in *T. tufae* occurred recently during the separation of *Tokorhabditis* and *Auanema*, perhaps as an adaptation associated with live birth. The HSNs demonstrate considerable innovation across nematode species, shifting in anatomical position as well as their expression of molecular markers like serotonin in nematodes such as *Pristionchus pacificus* (Loer et al., 2026; Loer & Rivard, 2007; Ramadan et al., 2026).

Additionally, *T. tufae* possesses sphincter-like vulval muscles that do not resemble typical nematode egg-laying muscles. Similar rearrangement of egg-laying muscles into sphincter-like muscles have been observed in some live-bearing flies (Meier et al., 1999). These observations highlight the remodeling of an ancestral egg-laying circuit in *T. tufae* and represent innovations in reproductive structure associated with its live-bearing life cycle.

Retention and internal hatching of eggs can be adaptive on its own. In *C. elegans*, natural genetic variation can result in internal hatching of eggs, live birth, and subsequent protection of progeny from environmental stressors such as osmotic and acidic stress (Mignerot et al., 2024; Vigne et al., 2021). However, this protection is at the cost of the mother’s fecundity and survival, as internal larvae can kill the mother. Other live-bearing nematodes like *Panagrellus redivivus* may avoid matricide by releasing early-stage larvae soon after hatching (Hechler, 1970; Srinivasan et al., 2013). Larvae are quiescent in *T. tufae* during their development *in utero*; future studies can determine whether this behavior protects the mother.

### Protective advantages of live birth

Live birth of progeny that are provisioned with nutrients *in utero* (viviparity) has evolved once in mammals and frequently in invertebrates, fish, and reptiles (Blackburn, 1999; Ostrovsky et al., 2016; Whittington, 2021). Among vertebrates, the squamate reptiles have experienced over 100 independent transitions from egg laying to viviparity, which has been suggested to protect progeny from environmental threats such as harsh temperature and predation (Ma et al., 2018; Whittington et al., 2022). Among extremophiles, viviparity has been observed in poecilid fish that thrive in sulfidic springs, suggesting a protective role of viviparity against chemical extremes (Greenway et al., 2024; Riesch et al., 2014).

A correlation between viviparous life cycles and toxic environments is found in nematodes as well. One clue for the evolution of viviparity in nematodes comes from the fact that viviparous *Tokorhabditi*s species isolated in Japan and the USA occupy a specialized niche, co-habitation with dung beetles (Ikeda et al., 2024; Ragsdale et al., 2022). Dung beetles live in feces and carrion, environments that exert strong osmotic stress and which are rich in phosphate, a metalloid chemically similar to the most commonly occurring arsenic ions. Thus, viviparity in *Tokorhabditis* may have evolved to support larval survival under osmotic and chemical stress. These observations across clades suggest that viviparity may be an evolvable motif that supports adaptation to stressful chemical extremes.

## Materials and Methods

### Animal strains and cultivation

*Caenorhabditis elegans* PD1074 (Yoshimura et al., 2019), *Auanema freiburgense* SB372, *Auanema rhodense* SB347 (Kanzaki, Kiontke, et al., 2017; Sudhaus, 2023), *Tokorhabditis tufae* PS8402 (Kanzaki et al., 2021; Shih, Lee, Shinya, et al., 2019), and *Tokorhabditis atripennis* NKZ329 (Ragsdale et al., 2022) were used as wild-type strains. *C. elegans* CB1372 *daf-7*(*e1372*), ECA581 *dbt-1*(C78S) (Zdraljevic et al., 2019), and the Unc mutant *T. tufae* CX1212 *ky1212* were used as mutant strains.

All animals were maintained at room temperature (20-23°C) using standard *C. elegans* culturing protocols (Brenner, 1974) on 6 cm plates containing Nematode Growth Media (NGM: 3 g NaCl, 2.5 g peptone, 22 g agar, 1 mL cholesterol (5 mg/mL in ethanol), 975 mL distilled H_2_O, autoclaved followed by adding 1 mL of 1 M CaCl_2_, 1 mL of 1 M MgSO_4_, 3.3 mL of 1 M K_2_HPO_4_, and 21.7 mL of 1 M KH_2_PO_4_ buffer sequentially) and *Escherichia coli* strain OP50 as a food source.

Unless otherwise indicated, experiments were performed using day-2 adults, which are at the peak of reproductive productivity in *T. tufae* and *C. elegans* (Kanzaki et al., 2021; Scharf et al., 2021). Synchronous populations of day-2 adults were prepared by picking L4 larvae and incubating them at 20°C for two days prior to testing.

Developmental stages in *T. tufae* were determined by the following methodology: Eggs, L1s, L2s, and dauers were obtained by dissecting adult animals. Eggs did not survive outside of the mother and began to swell and disintegrate 5 minutes after dissection, consistent with (Yamashita et al., 2023). A subset of dissected larvae also did not survive outside of the mother (**Supplemental Figure 1E-G**); these larvae were presumed to be L1s due to their small size compared to larvae that survived dissection (**Supplemental Figure 1H**). The larvae that survived dissection included L2s and dauers (**Supplemental Figure 1E-H**). Dauers were defined by their cuticle specializations and SDS resistance (**Figure 1C** and **Supplemental Figure 1I-J**). Surviving larvae were never observed to molt into L2, consistent with inviability of the L1s. Larvae that survived dissection but did not possess dauer-like characteristics were designated as L2s. In agreement with this assignment, they directly molted into the dauer form. Molting L2 larvae were identified by the presence of loose, separated cuticle and, in some cases, the early formation of faint dauer alae. The first stage after the dauer molt was identified as the L4 stage.

The timing of *T. tufae* development *in utero* was determined using an Olympus BX53 differential interference contrast microscope equipped with a Hamamatsu ORCA-spark camera. Observations of fertilization, hatching, and embryo growth *in utero* were obtained by paralyzing hermaphroditic adults with 0.1 M sodium azide in M9 buffer and mounting on 2% agarose pads followed by microscope observation. The timing of molts was determined by incubating nematodes at 20°C on NGM and recording the presence of shed cuticles on the culture plate at two-hour intervals (**Supplemental Figure 2**).

Mating crosses in *T. tufae* were performed by collecting the rare males and females that arise in every generation (Kanzaki et al., 2021). Females were collected by transferring L4 animals and incubating at 20°C for 2 days, after which day-2 adult females could be distinguished from hermaphrodites by their smaller size and accumulation of unfertilized oocytes. Males were collected by their distinctive tail morphology. Crosses were generated by transferring females and males, in a 1:3 to 1:6 ratio, to NGM plates with a 1 μL lawn of OP50 and incubated at 20°C for 3 days.

### Dissections

*T. tufae* adults on their second day of adulthood were dissected using 21-gauge and 30-gauge hypodermic needles (BD PrecisionGlide 305167 and 305106). Animals were bisected anterior to the anus or posterior to the pharynx to release the uterus. When needed, additional cuts were made to the uterus to release its contents of embryos and larvae. Embryos and L1 larvae that were not viable outside of the uterus (Yamashita et al., 2023 and **Supplemental Figure 1E-H**) were excluded from subsequent analysis. In assays comparing dissected larvae to dauer larvae, newly emerged dauers were obtained by transferring *T. tufae* adults at their second day of adulthood to NGM plates seeded with OP50, incubating at room temperature for 2 hours, and collecting the progeny that emerged.

### Survival assays

Survival in SDS was assessed by exposing animals in microtiter wells to 1% SDS (Millipore Sigma L4390) diluted in M9 and scoring for survival after 30 minutes (Cassada & Russell, 1975; Shih, Lee, & Sternberg, 2019).

Dose response arsenic toxicity assays were performed as in (Liao & Yu, 2005). Per assay, 20 animals were treated in 24-well culture plates (Corning CLS353047) with 0, 10, 20, 30, 40, 50, or 60 mM sodium arsenate dibasic heptahydrate (As(V), Millipore Sigma S9663) in K-media (53 mM NaCl and 32 mM KCl in sterile water). Animals were incubated at 20°C for 24 hours before scoring for survival.

Acute arsenic survival assays were performed as in (Shih, Lee, Shinya, et al., 2019). Per assay, 15 animals were treated in 24-well culture plates to 30 mM As(V) in sterile water. Animals were incubated at 20°C and scored for survival at 1, 2.5, 5, and 7 hour time points.

Survival in Mono Lake salinity was assessed using artificial Mono Lake water (MLW) made by dissolving 0.65 M NaCl, 0.1 M Na_2_SO_4_, 0.02 M K_2_SO_4_, 0.9 mM K_3_PO_4_, 0.175 M NaHCO_3_, 0.22 M Na_2_CO_3_, 0.045 M H_3_BO_3_ in sterile water (94.4 g/L final salinity) to approximate the chemical composition of natural Mono Lake water (Jones & Stokes Associates, 1993; Van Breugel & Dickinson, 2017). Artificial MLW was diluted in sterile water to the salinity of the sediments from which Mono Lake nematodes including *T. tufae* were isolated (Shih, Lee, Shinya, et al., 2019). Dose response and acute survival assays in dilutions of MLW were performed as above.

Population growth of animals was measured by cultivating ten day-2 adults on agar plates with NGM, Mono Lake salinity, or arsenic. Mono Lake salt plates were made by replacing the salts in NGM with 5.2 g/L artificial Mono Lake water; arsenic plates were made by diluting arsenic stock solution in NGM to a final concentration of 5 mM As(V). Plates were seeded with 100 µL of heat-killed OP50 (100 mg/mL). Growth was assessed by counting the number of viable progeny produced after 48 hours, which limited progeny production to a single generation.

In all survival assays, animals were scored as dead if they exhibited at least two of the following characteristics: absence of spontaneous or touch-evoked swimming, gray discoloration, tissue degradation, rupturing of the body wall, or a straight and rigid posture.

The salinity and ionic composition of all solutions used in survival assays are listed in Supplemental Table 1.

### Pharmacology

Pharmacological assays using exogenous serotonin, levamisole, and tyramine followed methods adapted from (Collins et al., 2016; Trent et al., 1983). Solutions of serotonin (Millipore Sigma H7752) and levamisole (Millipore Sigma L9756) were prepared by dissolving in M9 buffer, while tyramine (Millipore Sigma T2879) was dissolved in NGM plates. Birth rate measurements (described in further detail below in **Behavioral analysis**) were performed by exposing four day-2 adults to serotonin or levamisole in microtiter wells, or to tyramine in NGM plates seeded with OP50 bacteria. Progeny were counted after 3 hours for serotonin and levamisole experiments, and after 1 hour (*C.* elegans) or 2 hours (*T. tufae*) for tyramine experiments. Locomotor paralysis by exposure to exogenous serotonin was measured after 1 hour as previously described (Gürel et al., 2012; Ishita et al., 2021).

### Immunostaining

Animals were fixed and stained with DAPI (Millipore Sigma D9542), anti-serotonin antibody (Millipore Sigma S5545), and Acti-stain 488 Phalloidin (Cytoskeleton Inc. PHDG1) following methods adapted from (Loer & Rivard, 2007). Mixed-stage populations of animals were washed off NGM plates and rinsed three times with M9 buffer, fixed overnight at 4°C in 4% paraformaldehyde/PBS, rinsed three times in 0.5% Trixon X-100/PBS, incubated overnight at 37°C in 5% β-mercaptoethanol/1% Triton X-100/0.1 M Tris (pH 7.4), rinsed twice in 1% Triton X-100/0.1 M Tris, rinsed in collagenase buffer (1 mM CaCl_2_/1% Triton X-100/0.1 M Tris), and incubated for 30 minutes to 4 hours in 500 Units/mL Collagenase type IV (Millipore Sigma C5138) in collagenase buffer to digest the animals’ cuticles to make their tissues accessible for staining. Following digestion, animals were rinsed three times in 0.5% Triton X-100/PBS, incubated for 1 hour at room temperature in 1% bovine serum albumin (BSA; Millipore Sigma A9306)/0.5% Triton X-100/PBS blocking solution, and incubated overnight at room temperature in 1:100 anti-serotonin serum and 1:100 Acti-stain 488 in 1% BSA/0.5% Triton X-100/PBS. Following staining, animals were rinsed two times in 0.5% Triton X-100/PBS, incubated for 1 hour in 0.1% BSA/0.5% Triton X-100/PBS, then incubated for 2 hours at 37°C in 1:200 TRITC-conjugated goat anti-rabbit IgG (Millipore Sigma T6778) in 1% BSA/0.5% Triton X-100/PBS, rinsed three times in 0.5% Triton X-100/PBS, rinsed for 40 minutes in 1:1000 DAPI in 0.1% BSA/0.5% Triton X-100/PBS, then rinsed for 20 minutes in 0.1% BSA/0.5% Triton X-100/PBS. Animals were resuspended in ProLong Gold Antifade Mountant (Invitrogen P36930) and imaged at 40x using a Zeiss Axio Imager Z2 widefield fluorescence microscope, Zeiss Inverted LSM 980 laser-scanning confocal microscope, and Evident Inverted FluoView FV4000 laser-scanning confocal microscope.

Hybridization Chain Reaction Fluorescence *In Situ* Hybridization (HCR-FISH) was performed as described in (Loer et al., 2026; Ramadan & Hobert, 2024) with minor modifications. HCR was used to visualize the expression of *T. tufae* homologs of *tph-1*, *unc-86*, *egl-3*, *ast-1*, *sem-4*, and *nlp-3*. Homologs were identified from a draft genome of *T. tufae* containing transcript-level annotation using reciprocal blast, synteny, and peptide sequence analysis. HCR probes were designed for these genes using the HCR Probe Maker software (Kuehn et al., 2022). The maximum number of possible probes per gene were designed, ranging from 5 to 36, and ordered as Oligo Pools from IDT (see **Supplemental Table 2** for gene and probe sequences). Hairpins conjugating B1, B2, B3, and B4 initiator sequences with Alexa Fluor (AF) 647, AF 546, AF 488, and AF 594, respectively, were obtained from Molecular Instruments, along with Probe Hybridization Buffer, Probe Wash Buffer, and Amplification Buffer. All centrifugation steps were performed at 1200 g for 2 minutes.

Animals were fixed for HCR staining by washing mixed stage populations off NGM plates, rinsing three times with M9 buffer, and flash-freezing in 4% paraformaldehyde/PBS at -80°C. For staining, animals were then thawed at room temperature for 15 minutes, washed twice with PBST, incubated with 100 µg/mL proteinase K at 37°C for 15 minutes, washed twice with PBST, treated with 2 mg/mL of glycine solution on ice for 15 minutes, washed twice with PBST, then washed in 50% PBST / 50% probe hybridization buffer, pre-hybridized in Probe Hybridization Buffer at 37°C for 1 hour, and hybridized with 2 pmol of probes per gene at 37°C overnight. Samples were then washed four times in Probe Wash Buffer at 37°C, washed twice in 5x SSCT at room temperature, pre-amplified with Amplification Buffer at room temperature for 30 minutes, and incubated with snap-cooled hairpins at room temperature overnight. For imaging, samples were washed three times in 5x SSCT at room temperature, incubated in 1:1000 DAPI in 5x SSCT at room temperature for 30 minutes, washed once in 5x SSCT at room temperature, resuspended in ProLong Gold Antifade Mountant, and imaged at 40x using a Zeiss Inverted LSM 980 and Evident Inverted FluoView FV4000.

### Behavioral analysis

Video recordings of *T. tufae* live birth and *C. elegans* egg laying were acquired using custom moving-stage microscopes that automatically track moving animals (Cermak et al., 2020). Day-2 adults were transferred to NGM buffer on 2% agarose pads and recorded under a 10x objective lens (Olympus UPLFLN 10X 2) with Blackfly (Teledyne FLIR BFLY-U3-23S6M-C) or Chameleon (Teledyne FLIR CM3-U3-13Y3M-CS) CMOS cameras at 20.0 or 120.0 frames per second, respectively. Recordings were captured using LabView software and analyzed using ImageJ.

Ethograms of *T. tufae* live birth and *ex utero* activity of dissected larvae were generated by transferring animals to NGM plates seeded with 10 μL OP50. Plates were illuminated by an LED back light (Metaphase FR-BL3.5X6-W-24-ILD-BC), recorded under a Nikon Micro-NIKKOR 55mm f/2.8 lens with Blackfly CMOS cameras at 3.0 frames per second using SpinView software, and analyzed using ImageJ.

Birth rate measurements were performed by transferring four day-2 adults to microtiter wells containing assay solutions diluted in M9 and counting the progeny born after three hours. Assays with day-5 hermaphrodites and females (**Figure 6E**) were performed with single adults per replicate due to the rare occurrence of *T. tufae* females, and progeny were counted after 24 hours.

Live birth in SDS-killed adults was assessed by exposing *T. tufae* day-2 adults to 0.3% SDS for four minutes to kill the adults. Successful killing of adults was determined by the absence of spontaneous or touch-evoked movement, gray discoloration, tissue degradation, straight and rigid posture, and presence of *in utero* larvae in the extremes of the mother’s head or tail.

Vulval muscle contractions before, during, and after birth events were measured by tracking the distance between the vulval lips at ± 0, 0.03, 0.3, 1, 2, and 4 seconds around birth using ImageJ.

*In utero* larval behaviors were manually tracked inside day-2 adults. Postural measurements were made by manually drawing splines from the head to tail using ImageJ, and tortuosity was calculated as (spline length) / (shortest distance between the head and tail).

Matricide after internal larval growth was inferred if animals exhibited at least two of the following characteristics: (1) presence of larvae in the extremes of the head or tail of the mother, (2) L4 or older progeny observed in the mother, and (3) damage to the mother indicated by absence of spontaneous or touch-evoked crawling, gray discoloration, tissue degradation, or rupturing of the body wall.

### EMS Mutagenesis

Ethyl methanesulfonate (EMS) was used to perform forward mutagenesis in *T. tufae* as in (Pires Da Silva, 2013). To obtain synchronized populations of *T. tufae* for mutagenesis, mixed stages of *T. tufae* were washed off NGM plates, resuspended in 1% SDS in M9 for 10 minutes to collect dauers, washed three times in M9, transferred to 10 cm seeded NGM plates, and incubated at 20°C for 24 hours to allow dauers to develop into L4 larvae. L4s were transferred to a 15 mL tube, treated with 47 mM EMS at 20°C for 4 hours, washed three times with M9, then transferred onto 10 cm seeded NGM plates. P_0_ animals were grown for 3 days to generate F_1_ heterozygous progeny, washed in M9, resuspended in 1% SDS to collect F_1_ dauers, washed three times in M9, transferred to 10 cm seeded NGM plates, grown for 4 days to generate F_2_ progeny, washed in M9, resuspended in 1% SDS to collect F_2_ dauers, washed three times in M9, transferred to 10 cm seeded NGM plates at a density of 100 animals per plate, and grown for 2 days to generate young adults for phenotyping.

Phenotypes were assigned to *T. tufae* mutants by reference to known *C. elegans* mutants (Fay, 2013). *ky1212* was assigned a kinker Uncoordinated (Unc) phenotype as it demonstrated poor forward and backward movement with sharply angled body bends.

## Supporting information

Supplemental Materials (Figures, Tables, Video Legends, References)

## Acknowledgements

We thank the Rockefeller University and the Marine Biological Laboratory for generously providing facilities, resources, and support for this work. We thank M.B.H. Alfonso from the Rockefeller University’s Bio-Imaging Resource Center, RRID:SCR_017791, for help with imaging. We thank A. Pires-daSilva for *Auanema* strains and O. Hobert for reagents, and A. Sylvester, A. Michaiel, A. Bernard, E. Diaz, O. M. Arenas, B. Bertochi, T. Anttonen, I. Candido-Ferreira, D. Cortes, H. Farrants, T. Honda, I. Gonzalez Ishida, K. Jokura, M. Rodriguez-Santiago, S. Caetano Avritzer, A. Baker, C, Braendle, F. Buck, G. Donaldson, M. Ebert, Y. Greenfeld, H. Harris, O. Hobert, S. James, E. Jorgensen, P. Kidd, A. Kumar, L. Mignerot, E. New, M. Parent, J. Parker, D. Raizen, P.-Y. Shih, P. Sternberg, and R. Walsh for insightful comments. This work was initiated through a Kavli-Grass Fellowship supported by The Kavli Foundation to J.S.L. and supported by a grant from the Chan Zuckerberg Initiative to C.I.B.; a grant from the Human Frontier Science Program to C.I.B., R.S., and T.K.; and a Whitman Fellowship supported by competitive fellowship funds from Kuffler Research Awards, Robert Day Allen Research Awards, Baxter Post Doc Fellowship, L. & A. Colwin Summer Research Fellowship Fund of the Marine Biological Laboratory in Woods Hole, MA to J.S.L.

## Author Contributions

Conceptualization, J.S.L. and C.I.B.; Formal analysis, J.S.L., R.S., T.K., and C.I.B.; Investigation, J.S.L., S.O., C.G., H.Y., T.Y., R.S., and T.K.; Resources, J.S.L., R.S., T.K., and C.I.B.; Writing – original draft, J.S.L. and C.I.B.; Writing – review and editing, J.S.L., S.O., C.G., H.Y., T.Y., R.S., T.K., and C.I.B.; Visualization, J.S.L., H.Y., T.Y., and C.I.B.; Funding Acquisition, J.S.L., R.S., T.K., and C.I.B.

