## Supplemental Materials (Figures, Tables, Video Legends, References) for "Live birth protects the developing progeny of the extremophile nematode *Tokorhabditis tufae*"


- Supplemental Figures 1-8 (pages 2-9)
- Supplemental Tables 1-2 (pages 10-14)
- Supplemental Video Legends (page 15)
- Supplemental References (page 16)

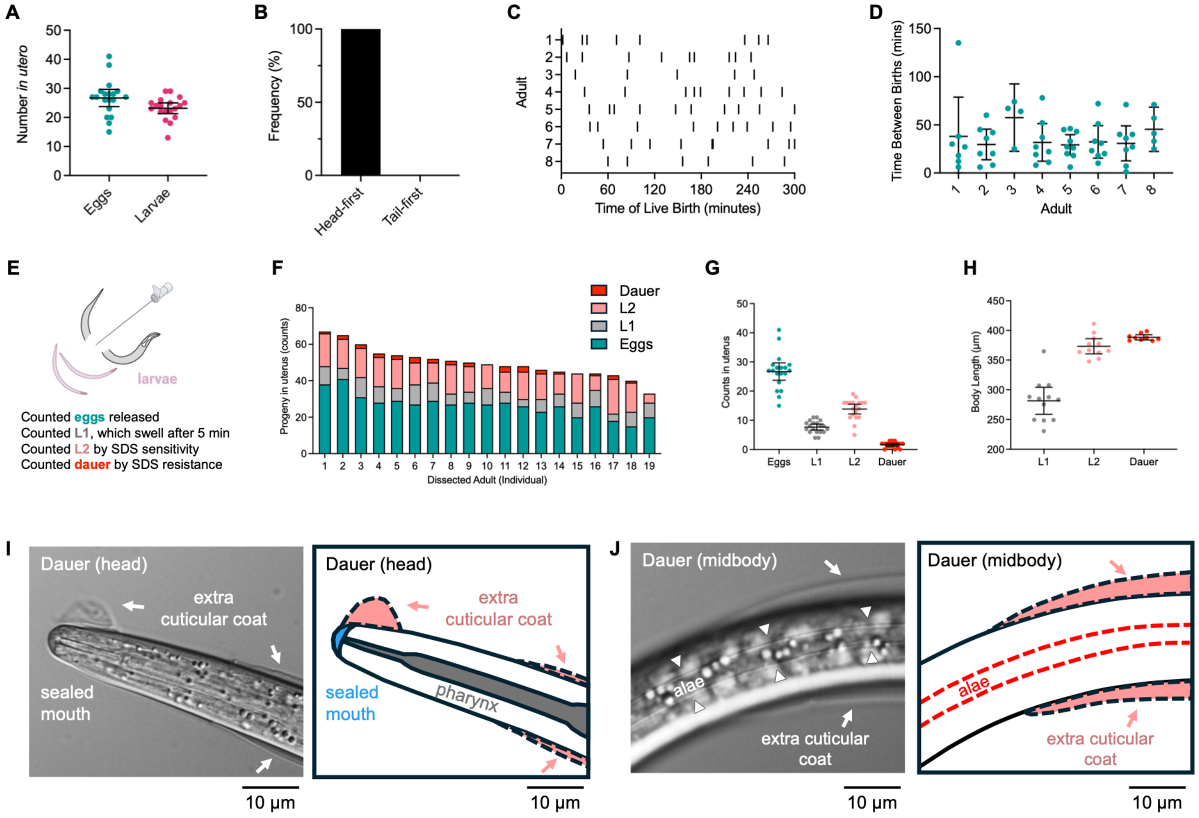


**Supplemental Figure 1. *Tokorhabditis tufae* progeny emerge at the dauer larval stage. (A)** *T. tufae* mothers hold an average of 26 eggs and 23 larvae *in utero*. n = 19 dissected adults. Error bars indicate 95% confidence interval. **(B)** *T. tufae* larvae emerge head-first during live birth (n = 40 events from 17 adults). **(C-D)** *T. tufae* mothers give birth once every 36.8 minutes, on average. (C) Ethogram of live birth events and (D) time between birth events (n = 65 births, 8 mothers). Error bars indicate 95% confidence interval. **(E-H)** Eggs, L1, L2, and dauer larvae *in utero*. (E) Schematic of methods used to characterize *in utero* progeny by their stage-specific features. (F) Counts and (G) summary statistics of *in utero* eggs, L1, L2, and dauers (dissected from 19 adults). (H) Body lengths of L1, L2, and dauer larvae (n = 9-12). Error bars indicate 95% confidence interval. **(I-J)** Newly born *T. tufae* larvae possess morphological features typical of nematode dauer larvae. Left: micrograph; Right: schematic highlighting dauer features in the micrograph. (I) Head of a dauer larva, showing a sealed mouth and ensheathing L2 cuticle. (J) Midbody of a dauer larva, showing distinctive alae.

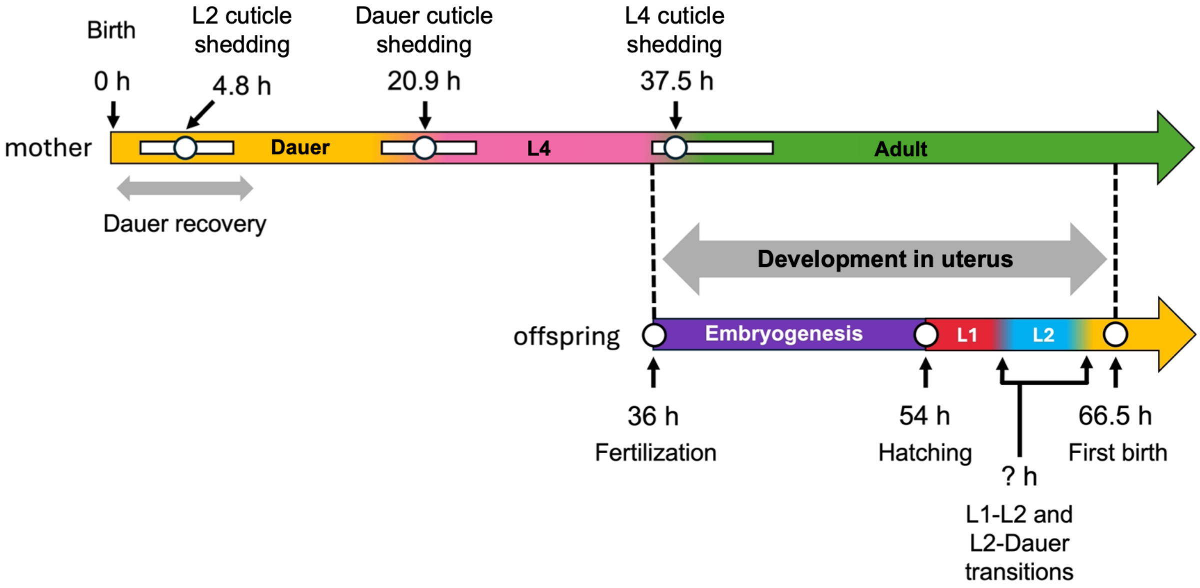


**Supplemental Figure 2. Timeline of developmental events in *T. tufae* mothers and offspring.** *T. tufae* dauers are born retaining their L2 cuticle as a sheath over their dauer cuticle, similar to dauers in *Auanema* (Kanzaki et al., 2017). In favorable conditions, they then initiate dauer recovery, defined by shedding of the L2 cuticle, opening of the mouth, and resumption of gonadal development. The timing of this event was determined from observations of 42 animals. The timing of subsequent cuticle shedding events was determined from 42 animals (dauer cuticle shedding) to 69 animals (L4 cuticle shedding). The timing of fertilization, hatching, and first birth were observed from 5-33 adult hermaphrodites. The timing of presumed L1-to-L2 and L2-to-dauer transitions during *in utero* development are unknown.

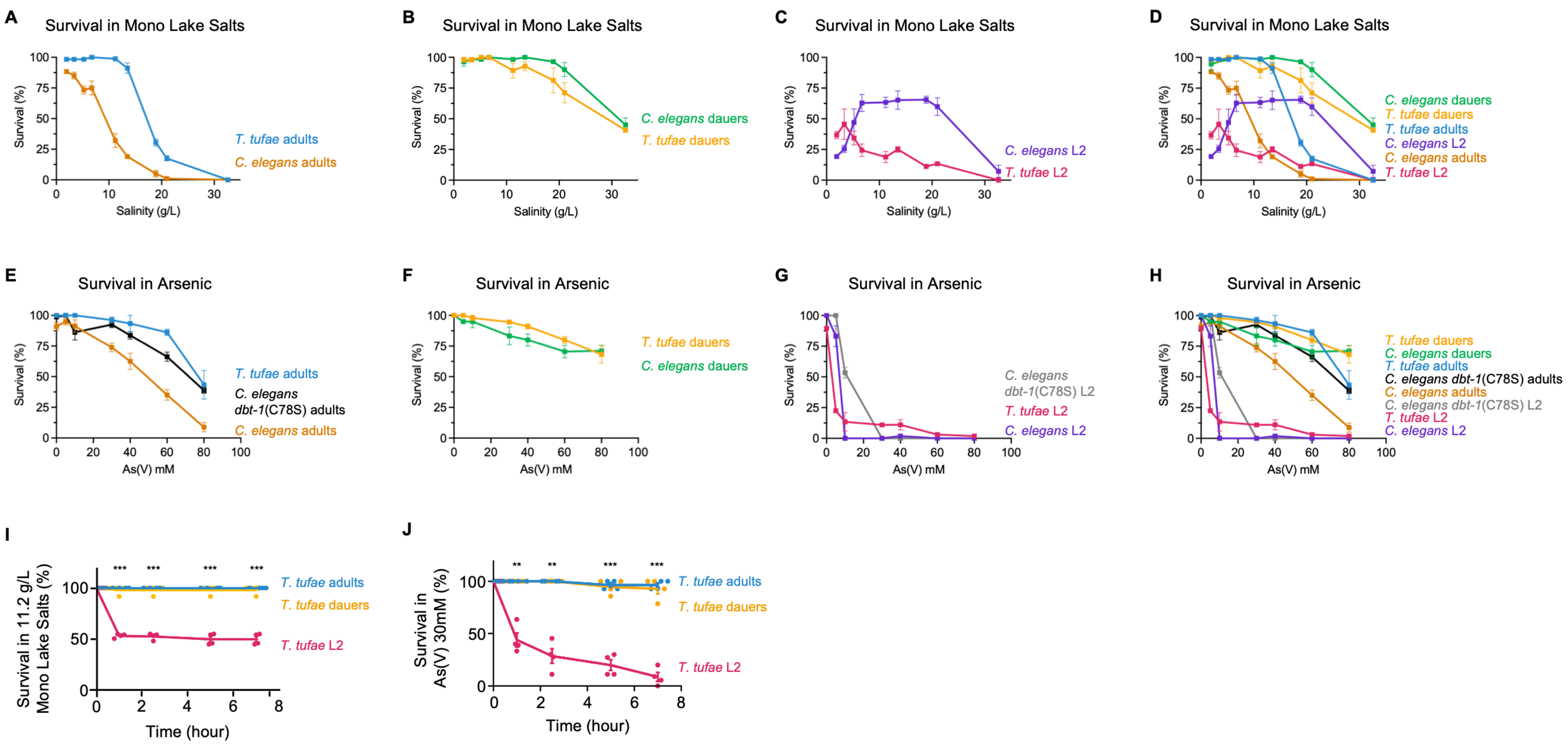
A
Survival in Mono Lake Salts
Survival (%)
Salinity (g/L)
T. tufae adults
C. elegans adults
B
Survival in Mono Lake Salts
Survival (%)
Salinity (g/L)
C. elegans dauers
T. tufae dauers
C
Survival in Mono Lake Salts
Survival (%)
Salinity (g/L)
C. elegans L2
T. tufae L2
D
Survival in Mono Lake Salts
Survival (%)
Salinity (g/L)
C. elegans dauers
T. tufae dauers
T. tufae adults
C. elegans L2
C. elegans adults
T. tufae L2
E
Survival in Arsenic
Survival (%)
As(V) mM
T. tufae adults
C. elegans dbt-1(C78S) adults
C. elegans adults
F
Survival in Arsenic
Survival (%)
As(V) mM
T. tufae dauers
C. elegans dauers
G
Survival in Arsenic
Survival (%)
As(V) mM
C. elegans dbt-1(C78S) L2
T. tufae L2
C. elegans L2
H
Survival in Arsenic
Survival (%)
As(V) mM
T. tufae dauers
C. elegans dauers
T. tufae adults
C. elegans dbt-1(C78S) adults
C. elegans adults
C. elegans dbt-1(C78S) L2
T. tufae L2
C. elegans L2
I
Survival in 11.2 g/L Mono Lake Salts (%)
Time (hour)
T. tufae adults
T. tufae dauers
T. tufae L2
J
Survival in As(V) 30mM (%)
Time (hour)
T. tufae adults
T. tufae dauers
T. tufae L2

**Supplemental Figure 3. Sensitivity of *T. tufae* and *C. elegans* to Mono Lake toxins. (A-D)** Survival of *T. tufae* and *C. elegans* after 24-hour exposure to Mono Lake salts. **(E-H)** Survival of *T. tufae* and *C. elegans* after 24-hour exposure to arsenic. **(I-J)** Acute survival of *T. tufae* adults, dauers, and dissected L2 in (I) 11.2 g/L Mono Lake salts and (J) 30 mM arsenic. For (A-H) n = 3-6 assays (10-45 animals per assay). Error bars indicate mean and SEM; (I-J) n = 4 assays (9-21 animals per assay). Error bars indicate mean and SEM. Statistics: (I-J) Two-way ANOVA with Tukey's multiple comparisons test. $^{**}p \leq 0.01$, $^{***}p \leq 0.001$ comparisons between *T. tufae* adults and dissected L2.

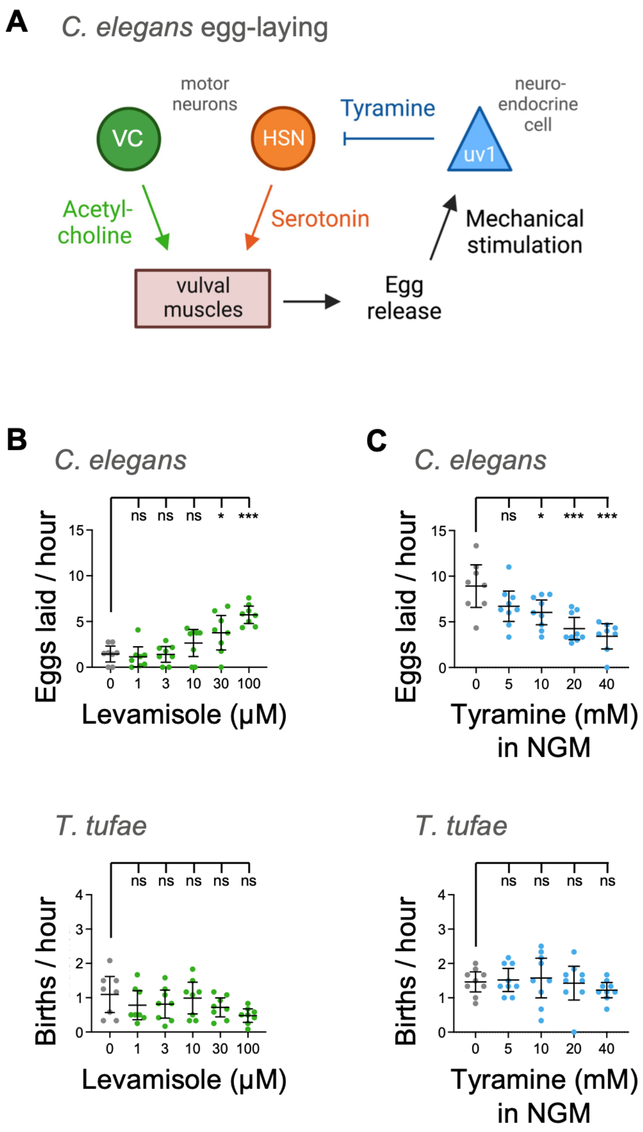


**Supplemental Figure 4. Canonical egg-laying circuits do not drive *T. tufae* live birth. (A)** Schematic of the *C. elegans* egg-laying circuit. **(B)** Levamisole, an acetylcholine agonist, increases egg-laying in *C. elegans* but does not affect birth rates in *T. tufae*. Levamisole is administered in M9, which inhibits basal egg laying in *C. elegans*. **(C)** Tyramine decreases egg-laying in *C. elegans* but does not affect birth rates in *T. tufae*. Tyramine is administered on NGM plates seeded with OP50. For each condition n $\geq$ 7 assays (4 animals per assay). Error bars indicate 95% confidence interval. Statistics: One-way ANOVA with Dunnett's multiple comparisons test. *p ≤ 0.05; ***p ≤ 0.001.

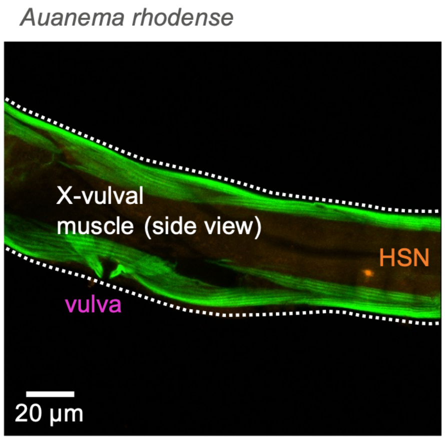


**Supplemental Figure 5. *Auanema rhodense* vulval muscles.** Antibody staining showing muscles (actin, green) and serotonin (orange) in the midbody of the egg-laying species *A. rhodense*. The animal is oriented anterior to the left and ventral down. Image is representative of n = 13 stained adult animals.

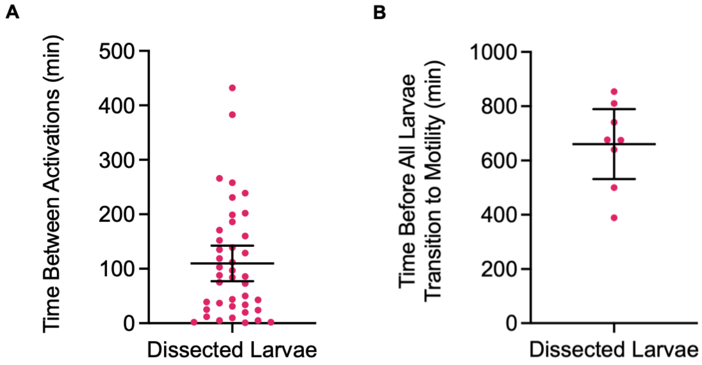


**Supplemental Figure 6. Dissected *T. tufae* larvae recapitulate the transition from quiescence to activity. (A-B)** Quantification of quiescence-to-activity transition events shown in Figure 5D. (A) Duration of time between quiescence-to-activity transition events (n = 49). Intervals between transition events are longer than intervals between live births (Supplemental Figure 1C-D), likely reflecting differences between *ex utero* and *in utero* environments. (B) Total time elapsed before all larvae from eight dissected adults became active.

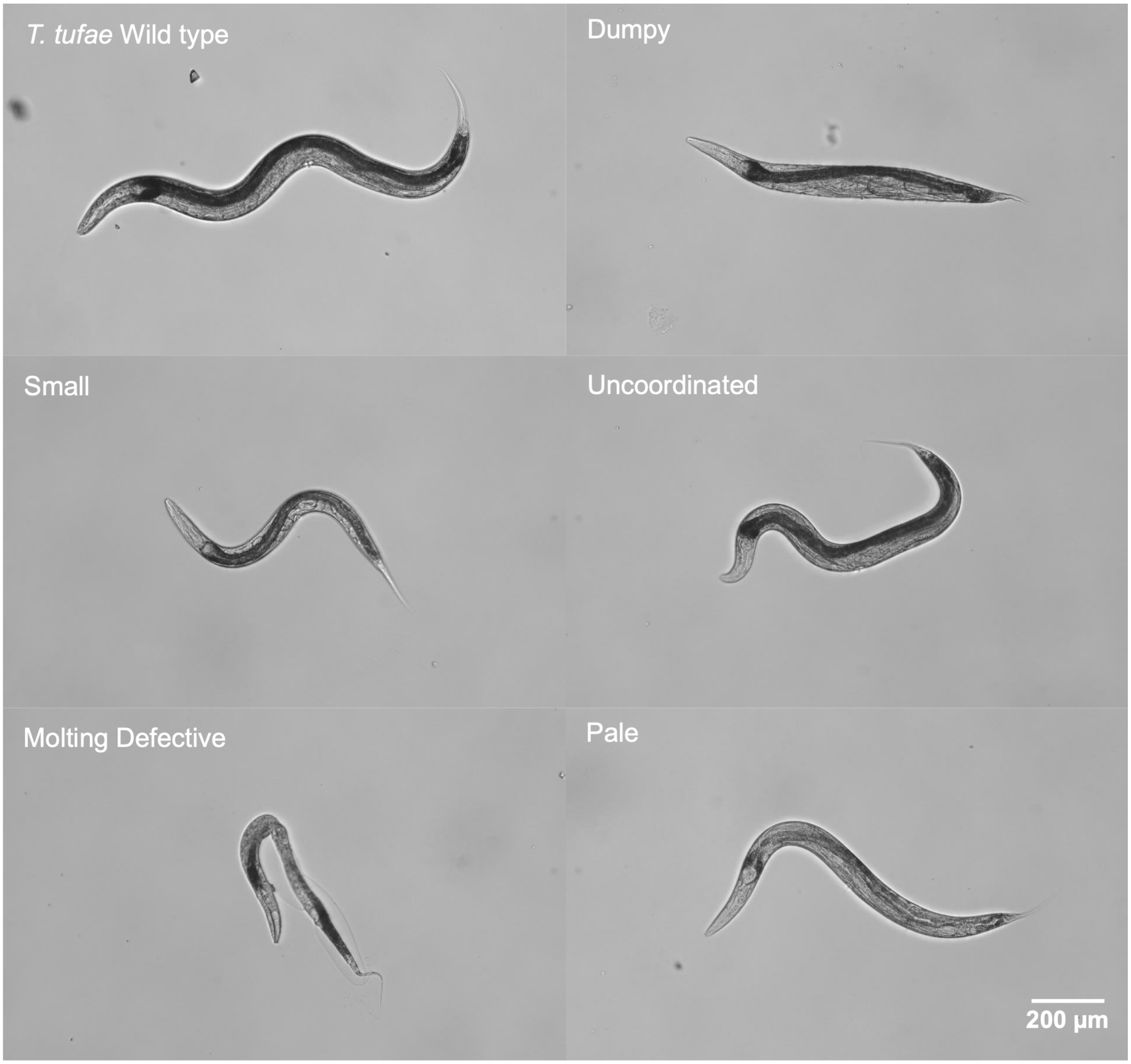


**Supplemental Figure 7. Ethyl methanesulfonate (EMS)-generated mutants of *T. tufae*.** Micrographs of *T. tufae* and mutants generated by EMS forward mutagenesis. All animals are oriented anterior left and ventral down; scale bar is 200 μm.

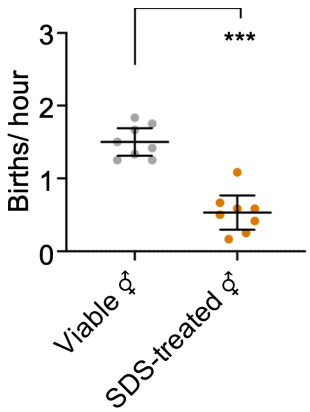


**Supplemental Figure 8. Birth rate of viable or SDS-treated adults.** n = 8 assays (4 animals per assay). Statistics: unpaired t test. ***p ≤ 0.001.

| Solution | Total Salinity (g/L) | pH | Carbonate (mM) | Calcium (mM) | Magnesium (mM) | Sodium (mM) | Potassium (mM) | Sulfate (mM) | Chloride (mM) | Boron (mM) | Phosphate (mM) | Arsenic (mM) |
|---|---|---|---|---|---|---|---|---|---|---|---|---|
| K-media | 5.5 | 4.5-5.0 | -- | -- | -- | 53.0 | 32.0 | -- | 85.0 | -- | -- | -- |
| Nematode Growth Media (NGM) | 6.7 | 5.5-6.0 | -- | 1.0 | 1.0 | 51.3 | 28.6 | 1.0 | 53.3 | -- | 25.2 | -- |
| M9 | 11.2 | 7.0 | -- | -- | 1.0 | 130.3 | 22.0 | 1.0 | 85.6 | -- | 44.4 | -- |
| S Basal | 12.8 | 6.0 | -- | -- | -- | 100.1 | 55.6 | -- | 100.1 | -- | 49.8 | -- |
| Mono Lake Water (MLW) (1.9 g/L) | 1.9 | 8.5-9.0 | 21.8 | -- | -- | 80.7 | 2.4 | 6.6 | 35.8 | 2.5 | 0.0 | -- |
| MLW (3.4 g/L) | 3.4 | 9.0-9.5 | 21.8 | -- | -- | 80.7 | 2.4 | 6.6 | 35.8 | 2.5 | 0.0 | -- |
| MLW (5.2 g/L) | 5.2 | 9.5 | 21.8 | -- | -- | 80.7 | 2.4 | 6.6 | 35.8 | 2.5 | 0.0 | -- |
| MLW (6.7 g/L) | 6.7 | 9.5-10.0 | 28.0 | -- | -- | 104.0 | 3.0 | 8.5 | 46.1 | 3.2 | 0.1 | -- |
| MLW (11.2 g/L) | 11.2 | 9.5-10.0 | 46.9 | -- | -- | 173.8 | 5.1 | 14.2 | 77.1 | 5.3 | 0.1 | -- |
| MLW (13.5 g/L) | 13.5 | 9.5-10.0 | 56.4 | -- | -- | 209.3 | 6.1 | 17.1 | 92.9 | 6.4 | 0.1 | -- |
| MLW (18.9 g/L) | 18.9 | 10.0 | 79.0 | -- | -- | 293.0 | 8.5 | 24.0 | 130.0 | 9.0 | 0.2 | -- |
| MLW (21.0 g/L) | 21.0 | 10.0 | 87.8 | -- | -- | 325.6 | 9.5 | 26.7 | 144.4 | 10.0 | 0.2 | -- |
| MLW (32.6 g/L) | 32.6 | 10.0 | 136.2 | -- | -- | 505.2 | 14.7 | 41.4 | 224.1 | 15.5 | 0.3 | -- |
| 5 mM As(V) in K-media | 6.4 | 8.0-8.5 | -- | -- | -- | 63.0 | 32.0 | -- | 85.0 | -- | -- | 5.0 |
| 10 mM As(V) in K-media | 7.3 | 8.0-8.5 | -- | -- | -- | 73.0 | 32.0 | -- | 85.0 | -- | -- | 10.0 |
| 20 mM As(V) in K-media | 9.2 | 8.0-8.5 | -- | -- | -- | 93.0 | 32.0 | -- | 85.0 | -- | -- | 20.0 |
| 30 mM As(V) in K-media | 11.0 | 8.5 | -- | -- | -- | 113.0 | 32.0 | -- | 85.0 | -- | -- | 30.0 |
| 40 mM As(V) in K-media | 12.9 | 8.5-9.0 | -- | -- | -- | 133.0 | 32.0 | -- | 85.0 | -- | -- | 40.0 |
| 50 mM As(V) in K-media | 14.7 | 8.5-9.0 | -- | -- | -- | 153.0 | 32.0 | -- | 85.0 | -- | -- | 50.0 |
| 60 mM As(V) in K-media | 16.6 | 9.0 | -- | -- | -- | 173.0 | 32.0 | -- | 85.0 | -- | -- | 60.0 |
| 80 mM As(V) in K-media | 20.3 | 9.0 | -- | -- | -- | 213.0 | 32.0 | -- | 85.0 | -- | -- | 80.0 |
| 5 mM As(V) in NGM | 7.7 | 6.0 | -- | 1.0 | 1.0 | 61.3 | 28.6 | 1.0 | 53.3 | -- | 25.2 | 5.0 |
| 30 mM As(V) in Water | 5.5 | 8.5-9.0 | -- | -- | -- | 60.0 | -- | -- | -- | -- | -- | 30.0 |

**Supplemental Table 1. Salinity and ionic composition of solutions used in study.** For reference, the salinity of freshwater is less than 0.5 g/L, brackish water is 0.5 – 30.0 g/L, and ocean water is 30.0 – 40.0 g/L (“The Venice System for the Classification of Marine Waters According to Salinity,” 1958).

| Gene | Coding Sequence |
| --- | --- |
| *Ttu_tph-1* | ATGGCATCTGCTATGAAGTTTCTTTATTATTCCAAAAAACCTCCTCACCGTACCTTGTCGAATACACTATCAAATGGCGACAGTCGGCTTGAAGATTTCAAACGACGGTTCCGCCGTAGTGGT<br>TCTCTGGGGATTCCGTTCGTACCAGAAGAGGACGTACACGACTTATTTTGCGATAAACTGGACTCACCTCTTCCAGATGAAGAAGGATATCCTATTTTAACTGTAATCATACGGTGCAACAG<br>GGCTGTCAACGGAATAGCCGAGATGGTTGGCAAACTTCCGGCTGATACATGCATTCGACATATGGAGAGCAGAGACTGTGAAGATGGTTCGACCGTCCACATGGACATTCTTGTTGAACTC<br>GAGCTACAGAAAGGCATGAATAACCAAGACGCTATGAAAATTATGCAACGCCAGGGACTCAATGTTTATGAAGTTTCGTTTAATATAATGCCATCAATCACAGACAACACTGAGAAGTTT<br>GTAGAGCCAGGATCTAATGATGCTATGACAGGGTCTCCATGGTTTCCTAAGAACATTTATGATCTTGATATATGTGCTAAAAGGGTTGTGATGTATGGAGCAGGATTAGATGCTGATCATCC<br>AGGTTTCAAAGATATGGAATATCGACAGAGACGGTTGATGTTTGCAGACGTTGCTTTAAATTATAAACACGGCGAGCCTGTTCCTCGCATTGATTATTCAGAAACAGAGAAAAAAACTTGG<br>GGTATTATATATAAGAAGCTGCGTGAACTTCACAAAAAATATGCTTGTCAAGAATTCCTTGATAATTTTGAACTGTTGGAGCGTCATTGTGGCTATTCAGAAAACAATATACCGCAGTTGGA<br>AGACATTTGTCGTTTTCTAAAAGCTAAAACTGGGTTTCGTGTCCGACCAGTAGCAGGATATTTGTCAGCTCGTGATTTTTTGGCTGGATTGGCTTATCGTGTTTTTTTCTGCACTCAATATGTAC<br>GACATCATGCTGATCCACTCTATACACCCGAACCAGATACTGTGCATGAACTTATGGGCCATATGGCTTTGTTTGCTGATCCGGATTTCGCACAGTTCTCACAAGAAATAGGTCTGGCCAGTC<br>TGGGCGCAAGTGAAGATGATTTGAAGAAATTAGCTACACTGTACTTTTTCTCTATAGAATTTGGCCTATCCTGTGAAGAGAACTCGGATGAAAAAAAAACAACGGAGAAGAAAACCTACA<br>AAGTTTATGGTGCTGGTCTGTTGAGTAGTGGTGGAGAACTTCAACATGCTGTTGAAGGTCCTTCTACTGTACTAAGATTTGATCCCGATAGGGTAATTCAGCAAGAATGTCTCATCACTACCT<br>ATCAAGAGGCCTACTTCTACACTCGAAATTTCGAAGAAGCTCAGCAGAAGCTGAGAATGTTCACAAGTAGTATGAAGCGCCCATTCATAGTACGCTATAATCCATACTCCGAGAGTGTGGA<br>GGTTCTCAATAACAAACGATCTCTTATGTCTGCAGTTAATACATTACGTTCAGATATAAATCTCCTTGCATCCGCTCTACACTATGTGATCTAA |
| *Ttu_unc-86* | ATGTCATCTGACGCTTCCGACGTCGATTTCCTTATATTTCTCGTCGTCGTCGTCGTCGTCTATAATTCATTTCTCTGCAGAATCTCTGAGGTCGTGTGTGTGTGTGTGTGTGTATGTGGACGCCTT<br>GAAAGCAGCACGACGAAAACGACGACTACAACAACCAGAAAGTCTATAGCCACACCTTATTCACAACCTGCCAAGATCGATCAAACTCGTGACGATCCATCTCAAGAAAACAGTCAGAT<br>TACAAACTCTAATCTGCCTCTTACTTCGATCATCTACCTACGAAGCCAAGAGGCTACTACCAGAAGTCGTGCTCCAGAAGCCGAGCCTTTCAGCTATATGCAGGCTCCGCTTTACCCTTCTCT<br>CAACAGTTTGCAAGGTCAACTAGGTGGATCATTAACAAGTTCTGGACGATACTCTCCCAACTCTTACAAGAATAGAATGCAACCGTTCAACAACTCCTTGTTTGGTACTTTCGATGACTCTCT<br>GTTTCAAGCAAGAGCAGCACAAGCAGCCGCCCTCGCAGATATTGATGTCAAAAATGTGAATCTAACTAATCAATTGATGAGACCTCAAGACATGTTCAGTAACTATCTCATGTCATCGGGT<br>CAAGACACAACAAACATTTATCAAGGGCTTCCGACGTCGTCTGAACCTTTTGACGCTTCAACCTTGATACCAACTAGCAATGCTACTGAACAAACTTCCCATGAAGGAATGACTCCATTGCA<br>GCAGAACTATGGTGCTCCATCCCATTTTACACAATACCCCATGACTCCATCTTTCGCTGTCTCAGCTCCCTCCCAAGTACGGTATCCACTATCAGCTCCCACTACTGAAATTGACACTGATCCC<br>AGAGAGTTGGAACGATTCGCCGAGCATTTCAAGCAACGACGTATCAAGCTAGGAGTCACGCAGGCGGATGTAGGTAAAGCATTAGCTCATCTCAAAATGCCAGGAGTTGGATCGTTATCC<br>CAATCAACAATTTGTCGATTTGAATCTTTAACCTTATCTCACAATAATATGGTGGCCTTGAAGCCAATTTTACATTCATGGCTTGAAAAGGCAGAAGAGGCTATGAAGCAGAAGGATTCCTG<br>TGGCGATGTATCTGGAATATTGCCAAACACAGACAAAAAACGGAAAAGAACAAGTATAGCTGCACCTGAAAAGAGACTGTTAGAAGACTATTTAGACAACAGCCACGTCCTTCGGGTG<br>AACGTATAGCACAAATAGCTGAGAAATTGGATTTGAAGAAAAACGTAGTCCGCGTTTGGTTCTGCAACCAACGACAGAAACAGAAAAGAGACTTCCGTTCACAATTCCGCTGTCGTAGCG<br>GCGGTATCAACGCTACCTCGACTCCTCGTCTAATGCCAACGACAACCAAATGTGCACCTTTTCCCGGATTACCAGGTTACTTTGAGTAG |
| *Ttu_egl-3* | ATGCAATGCCGGAATCCATTCTGGCTCAGCCTCTTTCTGGTAATCTTGTCTCTTACTGAGGCAGTAGAGGTTTTTACTAATCATTTTCATGTTCATGTAAAGGAACCCGGAATCGAGAATGCC<br>CATCGGATTGCTAAACGACACGGATTCATAAATAGAGGACCGGTGTTGGGAAGTGATACCGAATATCATTTTATTCAACCAGCTTTATCACATGCCAGAACTCGTCGATCAGTTGGACATC<br>ATGCCAAGCTCCACAAGGATAATGAGGTCAGTCACGTTGAGCAACTCACTGGATACAAACGAACAAAACGAGGATACAGACCTCTTGCCGAACGTCTTCAAACACAATTTGATTTCTCATC<br>CGTTCAATCACCCTCTGATCCTCTCTACCAATATCAATGGTATCTGAAGAACACTGGCCAAGCTGGAGGAAAAGCTCGTCTAGATCTTAATGTGGAGAAGGCATGGGCTCTCGGATTTACCG<br>GAAAAAACATTACCACCGCAATTATGGATGATGGTGTGGATTATATGCATCCAGATCTGAAGAATAACTTTAACGCTGAAGCCAGTTATGACTTCTCATCCAATGATCCATTCCCATACCCA<br>CGATACACCGATGACTGGTTCAACTCTCATGGAACTAGATGTGCTGGAGAAATTGCTGCAGCCAGAGACAACGGAGTCTGTGGAGTTGGAGTAGCTCCAGACAGCATGGCTAACTCCATG<br>GGACATGAGCCAAGCAAAATCCATATTTACTCTGCCAGCTGGGGACCAACCGATGACGGCAAGACTGTTGATGGACCTCGTAATGCTACAATGAGAGCTATTGTTAAGGGAGTCAATGAG<br>GGACGCGAAGGCTTGGGTTCCATCTTTGTTTGGGCTTCTGGTGACGGAGGAGAGGATGATGATTGCAATTGTGATGGTTATGCCGCATCGATGTGGACCATCTCCATCAATTCAGCCATCAA<br>CAATGGAGAAAATGCTCACTACGACGAGTCCTGCTCTTCAACTCTGGCTTCCACATTTTCAAACGGTGGTCGTAACCCAGAAACTGGAGTAGCTACCACTGACTTATACGGACGTTGCACTC<br>GTTCTCACTCTGGTACCTCAGCTGCTGCTCCGGAAGCAGCCGGAGTTTTTGCTCTTGCTCTCGAAGCTAATCCCAAGCTGACCTGGAGAGATCTCCAGCATTTGACTGTACTAACTTCTACAC<br>GTAACTCCTTGTTCGATGGCAGATGTCGTGATCTTCCAGAGCTCGGAATTGCTGAATATGATAACCACAAGAAGTCTACTGGCAACTGTACTCATTTCGAATGGCAGATGAACGGAGTTGGT<br>TTGGAATACAACCATTTGTTCGGATTCGGAGTTCTAGACGCTGCTGAAATGGTTATGCTGGCTATGGTCTGGAAAACTGCTCCACCCCGTTACCATTGTACTGCTGGAACGATCGATGTTCCT<br>CATGAGATTCCTGCTGACGGCAACCTCGTGCTCGATCTTGAAACCGATGGCTGTGCCGACTCAGCTACAGAGGTCCGATACTTGGAACATGCTCAAGCTGTTATCTCTTTCAACTCAACTCGT<br>CGTGGAGATACTACCCTCTACCTGATCTCACCCATGGGAACACGCACAATGATTCTTTCCCGACGTCCCAAAGATGATGATGCAAAGGATGGTTTCACTAACTGGCCATTCATGACCACCCA<br>CACTTGGGGAGAAAATCCAAGAGGAAAGTGGAGATTGGTTGCTCGATTCCAGGGACCCGGAGACCACAGAGGAACACTGAAGAAGTTCACTCTTATGCTCCACGGAACCAAGGAACCAC<br>CTTATTCGGGCATCGAGCCACTCATGGGACACGTCAACTCCAAGCTTCAGGTCGTTCAGACGGCTCATAAGCGTATGACTCATTAA |
| *Ttu_ast-1* | ATGACCTCCCTTAAACGGCTGTCAGCTATTGGAGTTAGCCGCTGGTCGGCTGTTATTGAATATGGTGAGTCTGTTAACATAGAGCAGCGATTTTCACCCAAAAGGGAAGGAAACGGATCCG<br>GAATGGACGTCAAGACGGAAAACGCCGCCACAGTTGCTAGCGCAGCCGTTGCTGTCAAAGCTGCTGAAGCAGTTTCCTTTCCATTTCTACAGAACGGTGTGCCCAATGGAAATCTTCCTCG<br>AGGTCTTTCCAGTTTTTATGAATCAAACAGTGGAACGATGGCTCAGACAACCAAAAACTCAGGTTCTGATGTCTTCTCGGGTATGAATCTCTCGATGCAAGCCTTCAACAGTTCGGCTGTTTC<br>TGCTACTGCTGCAGCAGCTGGAGCTGGAGCTAATCAAGTTAAAGATGAGACTCTTCAAGCTCCTCAATTCAACATGCCATGCTTCAACTATAACTTGAAGCATGTTATTCCAACCTGCAATC<br>CAAGCTCTTTCAGTACTCAAGATCCTTACCAGACTCTTTGTACTACCAGCAGACAATTGGCTCACTCAGGCAGTGGTCAGACTCAATTATGGCAATTCCTGTTGGAATTGCTGTCAGATGCTC<br>GTCATCAAGAAGTTATCACTTGGGAAGGAACTCAGGGCGAATTCAAACTTGTTGATCCGGATGAGGTTGCCAGAAAATGGGGAGAACGCAAATCGAAACCGAACATGAATTACGATAAG<br>ATGTCTCGTGCTCTAAGATATTATTATGACAAGAATATCATGGCCAAGGTTCATGGAAAACGATATGCTTACAAATTCGATTTTCAAGGAATTGCCGCTGCTATTCAACCTCAAACACCCAG<br>CTCAACGTCGGCAGATTATCTGGCTGTCAACCGATTACCTCAAGAATTCATGCAGGGTCCATGGAGCAATTATCGTTCTCTCATCCAACCCGGCTTCCAACCGGGTACAGCTCTCTTCACCCC<br>ATCAAGTATGGGATATGGTAGTTTTTCAACAGCCAACAATACATCATTACCTACTAGAAACTTTCCACTTTACGGCTCTAATATGAGCAATTACTCCAAATGTGCATAA |
| *Ttu_sem-4* | ATGGAAGGCGTGGACGAGTCGTCGTCGCCGATTGATCTGTCGACGACGACGGCAAGCTCGGCCACAGCCTTGATGGACAAAATGCTAGATGATAACGGAAAATCCTTGTTGCTCCAATCC<br>CTAGCATCCATAGCTACACCTCAGCAGATTTTAACATTGTGCGCTCAATTAGCAGCTCAACGACAGCAACATCAATCTTTATCGTCAACCGAGCAGGAGCATGAGGTGAGCATGATGGTCG<br>ATGATTCGACTGCTGATGTAACGCCTCCGGCAACACCAAAAAAGGATGTAATAATCTGTGCTGCAAGTGGATGTGGACAGTGCTTCAGCTCACAAGGTGCTCATCTGTGGCATTTGATGGA<br>AGAACATTCAACAGAAAAGCTATTACAATGCGACGAATGTGGACAACGTTTTAGCGATCGTGATCAGATTATTGAACATGAGTGTGAACGACAGATGCGTTCACGATCAGCTGGTGCTGA<br>ACGATCGACTGTCAACAGTCCTGATTCTGCTGATCGTGGATCTGTTCCAGCTCAAGCTGGACCAAGCTTTTCCATTGTTGAAGCTGAAGACGATCTTGCCTTGAGCAGCTCGATGGCTGATGA<br>GGATGGTACTGGCGCCATGCTCTTACCTTCCTCAGCCAGTTTGCTCAACCAGCTTTTTGCTCAACAGCATGTCTCCATCAATTCGTCATCCAACCTATCCACCAACAATCATAATCTACACAAT<br>CAGCAACATATTCAAAATGCGCAGCATCTTCAGCAACTTCAGCAGCTTCAACAGCTTCAAAATCAACAACAACAACATCTGCTGGGTCTCACAAATGGCTCACTGAAGTCTGACAATTACT<br>CCATCGGTCCTCAAATGGCTTCAGTAGCTGCAGCTGCTGGTCTTTCTGGCGACAATCAGACGCCAACCAGCGCCAGCGGTGCTCCTGTCACGCCCACGTCGGCAGCCATGTTTCCATTATCG<br>CTGAACCCATTCGGACATCCGTCTGCTCCACCCTTTTTCATGAGACCATTTGATGGTCCGAGAGTCGACATGTTTCCTCCTTCAGGTGCTGGTGGAGTTCGACCGGAAATCAACAATGAGGAC<br>GACTGGGAAGCTTTGATGGAGATTTCGACAACGGATGAAGCCGAGAAGATCCGTGCTTTAGTTGGGGACAAAGCTATGCCGACGACAGATCCCAATCAATGCATCCTCTGTCGACGAGTTT<br>TGTCGTGCAAGTCAGCACTTCAGATGCATTACAGAACACACACAGATGCCTCAGATGCCACCGTTGGGAAATCATTTGTCAATGGCTGGACTGCAGCCACAACAGCAATGTCCCATATGCC<br>AGAAGAGGTTCAGCATTTTGATAGAACTATTTGTAATTTTTCTTACAGATGTATGACACCTTCGCAACTTCAAGCGCATATGGCTGAACATCGACAAGCGCTAACAGGACGGATACCCAGTG<br>GAATGCCGCAGGGATCTCTTGGATCCGGCGCCTCTTTTCCTGTTGGCCTACCTCTCTTCCCATTCTTTCCTCAGTCCTTTTCTGGATCCGGTGAGCAGACCGGGGCTGGAGTAGGAGCTGGAAC<br>CCAACCTTTTAACCTTGCTCAGCTGCTTGGCAAGCAATTAGCCACCGCAGAGGAGATTCGGGTGAAGCCAACTACGAGTTTATCCACTACTGGAAGTCCTCCCCCCCCAACTTCTTCCAGCG<br>ATTTAGACGAGAAGAAAGATGATGTGGAAATGGTTGGAAGTCCGGAACAGAAGCCCCTTCAACCCGGCGATGCCAACTCTTCTATTTCTCCATCAGCCAGTAGTGAAGCCTCCCATGATCA<br>AACAGACGGTTCTTCTGTGAGCCGTTGTGAATCTGAAGGCGAGATGAAAACGTCTGCAGCCGATGGTGCTCTTCTGAACACCTCGACTGCCAGGACAAGCACCAGCAAATCCTCCTCTTCC<br>GACATCAAGATGGACTGTTCGTCTGCTGATGAATCTGTTTCCACTCCACGATCAGCTCCTTTAGCTCCCTCATCTCCTCGGCTGACAGCAGATAATCCATTAGAAGCAATGCAGAAGATGTG<br>GGCTGAAACTGAACCACCTCCACCTCGACAAATTCCCGTGTTATCAAAGCATCAATGTGGTGTCTGCTTCAAGCATTTCAGCTCAAGCTCAGCTCTCCAGATACACATGAGAACTCATACTG<br>GAGATAAGCCGTTTAAATGTGAGATGTGTGGTCGAGCCTTCACCACCCGAGGGAATCTGAAAGTTCATATGGGTACTCATATGTGGCAACAAAGCCCAAGTAGGCGTGGACGACGAATTT<br>TCGACGCTTCCCCACCGACGCCGAATCCTGTAGACGTCGTCGTGTCACCTATTAAAGGCTCTTCCGCATCTCTCAGCTTGCCTGGTGGACTTCAACCTCCCCTTGGCCTTTTTCCAGCTGGAGT<br>GCCCAATATGGAAATGATGATGCTTTTATGGAGGACGGTATGCTCTGTTTGTCAAAAAGTTTGTGCTAGCCCATCAGAACTGGAGCAACATTTGCGGGATCATCTGGAGACGTCACACGCC<br>GGTACGACGCCATCCTCAGCAGCTAGTCCGCAACCTACTACTGCTGCCGAATAA |
| *Ttu_nlp-3* | ATGGATTTCTCGAATTTCTGGCTTCCTAAGGCTCTTATTTCAGGTTCAAGATGGGGCCGTGGACGAAAAAAATATAGTAAGCTAGCACGAGATCCAAATATTTTGCCAGTGCCGGCCAATAT<br>GAAGCTAGTTTTCACTGCTCTTCTCATCCTAGGAGCCGTCTTCTGCATTGTTCAAGCTGAAGAGCGTATTAGTACCTTCTCAGAGAATGGCAAACCTCTTATGGGATATATGATCTCTGGAAA<br>TCCCGGAAGGCAAATCCTATTCTATCCTCGTGAAGCCACTTTTAACATGGAAAAAAGATATTTCGATAGTTTGGCTGGGCAGTCTCTTGGCAAATAG |

**Supplemental Table 2. Gene and probe sequences used in study.**

| HCR Probe Pool | Probe Sequence |
|---|---|
| B3_*Ttu_tph-1* | GTCCCTGCCTCTATATCTTTAAGGAGATTTATATCTGAACGTAAT |
| | GTCCCTGCCTCTATATCTTTTTATTGAGAACCTCCACACTCTCGG |
| | GTCCCTGCCTCTATATCTTTGCTTCATACTACTTGTGAACATTCT |
| | GTCCCTGCCTCTATATCTTTAGTGTAGAAGTAGGCCTCTTGATAG |
| | GTCCCTGCCTCTATATCTTTCTATCGGGATCAAATCTTAGTACAG |
| | GTCCCTGCCTCTATATCTTTCTCCACCACTACTCAACAGACCAGC |
| | GTCCCTGCCTCTATATCTTTTAGGCCAAATTCTATAGAGAAAAAG |
| | GTCCCTGCCTCTATATCTTTTCACTTGCGCCCAGACTGGCCAGAC |
| | GTCCCTGCCTCTATATCTTTGATCAGCAAACAAAGCCATATGGCC |
| | GTCCCTGCCTCTATATCTTTTGTATAGAGTGGATCAGCATGATGT |
| | GTCCCTGCCTCTATATCTTTCTACTGGTCGGACACGAAACCCAGT |
| | GTCCCTGCCTCTATATCTTTCAACTGCGGTATATTGTTTTCTGAA |
| | GTCCCTGCCTCTATATCTTTTTATCAAGGAATTCTTGACAAGCAT |
| | GTCCCTGCCTCTATATCTTTCGAGGAACAGGCTCGCCGTGTTTAT |
| | GTCCCTGCCTCTATATCTTTGTCTCTGTCGATATTCCATATCTTT |
| | GTCCCTGCCTCTATATCTTTTCCATACATCACAACCCTTTTAGCA |
| | GTCCCTGCCTCTATATCTTTAACCATGGAGACCCTGTCATAGCAT |
| | GTCCCTGCCTCTATATCTTTTGTTGTCTGTGATTGATGGCATTAT |
| | GTCCCTGCCTCTATATCTTTCTGGCGTTGCATAATTTTCATAGCG |
| | GTCCCTGCCTCTATATCTTTAGTTCAACAAGAATGTCCATGTGGA |
| | GTCCCTGCCTCTATATCTTTCCATATGTCGAATGCATGTATCAGC |
| | GTCCCTGCCTCTATATCTTTGTTGACAGCCCTGTTGCACCGTATG |
| | GTCCCTGCCTCTATATCTTTTCTGGAAGAGGTGAGTCCAGTTTAT |
| | GTCCCTGCCTCTATATCTTTGTACGAACGGAATCCCCAGAGAACC |
| | GTCCCTGCCTCTATATCTTTAAGCCGACTGTCGCCATTTGATAGT |
| | TTAGATCACATAGTGTAGAGCGGATTTCCACTCAACTTTAACCCG |
| | TTAACTGCAGACATAAGAGATCGTTTTCCACTCAACTTTAACCCG |
| | ATGGATTATAGCGTACTATGAATGGTTCCACTCAACTTTAACCCG |
| | CTTCTGCTGAGCTTCTTCGAAATTTTTCCACTCAACTTTAACCCG |
| | GTGATGAGACATTCTTGCTGAATTATTCCACTCAACTTTAACCCG |
| | AAGGACCTTCAACAGCATGTTGAAGTTCCACTCAACTTTAACCCG |
| | TTTTTCATCCGAGTTCTCTTCACAGTTCCACTCAACTTTAACCCG |
| | AGTGTAGCTAATTTCTTCAAATCATTTCCACTCAACTTTAACCCG |
| | TTTCTTGTGAGAACTGTGCGAAATCTTCCACTCAACTTTAACCCG |
| | AAGTTCATGCACAGTATCTGGTTCGTTCCACTCAACTTTAACCCG |
| | AAAAATCACGAGCTGACAAATATCCTTCCACTCAACTTTAACCCG |
| | AGCTTTTAGAAAACGACAAATGTCTTTCCACTCAACTTTAACCCG |
| | CCACAATGACGCTCCAACAGTTCAATTCCACTCAACTTTAACCCG |
| | TTTTTCTCTGTTTCTGAATAATCAATTCCACTCAACTTTAACCCG |
| | TTAAAGCAACGTCTGCAAACATCAATTCCACTCAACTTTAACCCG |
| | ACCTGGATGATCAGCATCTAATCCTTTCCACTCAACTTTAACCCG |
| | ATATCAAGATCATAAATGTTCTTAGTTCCACTCAACTTTAACCCG |
| | TAGATCCTGGCTCTACAAACTTCTCTTCCACTCAACTTTAACCCG |
| | AAACGAAACTTCATAAACATTGAGTTTCCACTCAACTTTAACCCG |
| | TGGTTATTCATGCCTTTCTGTAGCTTTCCACTCAACTTTAACCCG |
| | TCGAACCATCTTCACAGTCTCTGCTTTCCACTCAACTTTAACCCG |
| | AAGTTTGCCAACCATCTCGGCTATTTTCCACTCAACTTTAACCCG |
| | ACAGTTAAAATAGGATATCCTTCTTTTCCACTCAACTTTAACCCG |
| | AAAATAAGTCGTGTACGTCCTCTTCTTCCACTCAACTTTAACCCG |
| | ACGGCGGAACCGTCGTTTGAAATCTTTCCACTCAACTTTAACCCG |
| B2_*Ttu_unc-86* | CCTCGTAAATCCTCATCAAAAAAAGGTGCACATTTGGTTGTCGTT |
| | CCTCGTAAATCCTCATCAAAATACCGCCGCTACGACAGCGGAATT |
| | CCTCGTAAATCCTCATCAAAGTTGGTTGCAGAACCAAACGCGGAC |
| | CCTCGTAAATCCTCATCAAATATTTGTGCTATACGTTCACCCGAA |
| | CCTCGTAAATCCTCATCAAAAACAGTCTCTTTTCAGGTGCAGCTA |
| | CCTCGTAAATCCTCATCAAAGATACATCGCCACAGGAATCCTTCT |
| | CCTCGTAAATCCTCATCAAAATGAATGTAAAATTGGCTTCAAGGC |
| | CCTCGTAAATCCTCATCAAATTCAAATCGACAAATTGTTGATTGG |
| | CCTCGTAAATCCTCATCAAATGAGCTAATGCTTTACCTACATCCG |
| | CCTCGTAAATCCTCATCAAAGCTTGAAATGCTCGGCGAATCGTTC |
| | CCTCGTAAATCCTCATCAAAAGTGGGAGCTGATAGTGGATACCGT |
| | CCTCGTAAATCCTCATCAAAGGAGTCATGGGGTATTGTGTAAAAT |
| | CCTCGTAAATCCTCATCAAAGAGTCATTCCTTCATGGGAAGTTTG |
| | CCTCGTAAATCCTCATCAAATGAAGCGTCAAAAGGTTCAGACGAC |
| | CCTCGTAAATCCTCATCAAATCTTGACCCGATGACATGAGATAGT |
| | CCTCGTAAATCCTCATCAAAGATTAGTTAGATTCACATTTTTGAC |
| | CCTCGTAAATCCTCATCAAATGCTCTTGCTTGAAACAGAGAGTCA |
| | CCTCGTAAATCCTCATCAAAGGTTGCATTCTATTCTTGTAAGAGT |
| | CCTCGTAAATCCTCATCAAAATCCACCTAGTTGACCTTGCAAACT |
| | CCTCGTAAATCCTCATCAAAATAGCTGAAAGGCTCGGCTTCTGGA |
| | CCTCGTAAATCCTCATCAAACGTAGGTAGATGATCGAAGTAAGAG |
| | CCTCGTAAATCCTCATCAAACTTGAGATGGATCGTCACGAGTTTG |
| | CCTCGTAAATCCTCATCAAAGGCTATAGACTTTCTGGTTGTTGTA |
| | CCTCGTAAATCCTCATCAAACGTCCACATACACACACACACACAC |
| | CCTCGTAAATCCTCATCAAAAATTATAGACGACGACGACGACGAC |
| | CTACTCAAAGTAACCTGGTAATCCGAAATCATCCAGTAAACCGCC |
| | ATTAGACGAGGAGTCGAGGTAGCGTAAATCATCCAGTAAACCGCC |
| | AACGGAAGTCTCTTTTCTGTTTCTGAAATCATCCAGTAAACCGCC |
| | GTTTTTCTTCAAATCCAATTTCTCAAAATCATCCAGTAAACCGCC |
| | CGTGGCTGTTGTCTAAAATAGTCTTAAATCATCCAGTAAACCGCC |
| | TTTTTGTCTGTGTTTGGCAATATTCAAATCATCCAGTAAACCGCC |
| | TCATAGCCTCTTCTGCCTTTTCAAGAAATCATCCAGTAAACCGCC |
| | CATATTATTGTGAGATAAGGTTAAAAAATCATCCAGTAAACCGCC |
| | AACGATCCAACTCCTGGCATTTTGAAAATCATCCAGTAAACCGCC |
| | GCGTGACTCCTAGCTTGATACGTCGAAATCATCCAGTAAACCGCC |
| | CTCTCTGGGATCAGTGTCAATTTCAAAATCATCCAGTAAACCGCC |
| | TGGGAGGGAGCTGAGACAGCGAAAGAAATCATCCAGTAAACCGCC |
| | ATGGAGCACCATAGTTCTGCTGCAAAAATCATCCAGTAAACCGCC |
| | AGTAGCATTGCTAGTTGGTATCAAGAAATCATCCAGTAAACCGCC |
| | GGAAGCCCTTGATAAATGTTTGTTGAAATCATCCAGTAAACCGCC |
| | TGAACATGTCTTGAGGTCTCATCAAAAATCATCCAGTAAACCGCC |
| | AATATCTGCGAGGGCGGCTGCTTGTAAATCATCCAGTAAACCGCC |
| | AAAGTACCAAACAAGGAGTTGTTGAAAATCATCCAGTAAACCGCC |
| | GAGAGTATCGTCCAGAACTTGTTAAAAATCATCCAGTAAACCGCC |
| | GAGAGAAGGGTAAAGCGGAGCCTGCAAATCATCCAGTAAACCGCC |
| | CGACTTCTGGTAGTAGCCTCTTGGCAAATCATCCAGTAAACCGCC |
| | GATTAGAGTTTGTAATCTGACTGTTAAATCATCCAGTAAACCGCC |
| | GATCTTGGCAGGTTGTGAATAAGGTAAATCATCCAGTAAACCGCC |
| | GTCGTTTTCGTCGTGCTGCTTTCAAAAATCATCCAGTAAACCGCC |
| | CGACCTCAGAGATTCTGCAGAGAAAAAATCATCCAGTAAACCGCC |

**Supplemental Table 2 continued. Gene and probe sequences used in study.**

| | |
|---|---|
| **B1_*Ttu_egl-3*** | GAGGAGGGCAGCAAACGGAACTGAACGACCTGAAGCTTGGAGTTG |
| | GAGGAGGGCAGCAAACGGAATAAGGTGGTTCCTTGGTTCCGTGGA |
| | GAGGAGGGCAGCAAACGGAATGTGGTCTCCGGGTCCCTGGAATCG |
| | GAGGAGGGCAGCAAACGGAATTCTCCCCAAGTGTGGGTGGTCATG |
| | GAGGAGGGCAGCAAACGGAATCATCATCTTTGGGACGTCGGGAAA |
| | GAGGAGGGCAGCAAACGGAATCAGGTAGAGGGTAGTATCTCCACG |
| | GAGGAGGGCAGCAAACGGAAAGCATGTTCCAAGTATCGGACCTCT |
| | GAGGAGGGCAGCAAACGGAAAGATCGAGCACGAGGTTGCCGTCAG |
| | GAGGAGGGCAGCAAACGGAACAGCAGTACAATGGTAACGGGGTGG |
| | GAGGAGGGCAGCAAACGGAACATTTCAGCAGCGTCTAGAACTCCG |
| | GAGGAGGGCAGCAAACGGAAACTCCGTTCATCTGCCATTCGAAAT |
| | GAGGAGGGCAGCAAACGGAATATCATATTCAGCAATTCCGAGCTC |
| | GAGGAGGGCAGCAAACGGAAGGAGTTACGTGTAGAAGTTAGTACA |
| | GAGGAGGGCAGCAAACGGAATTGGGATTAGCTTCGAGAGCAAGAG |
| | GAGGAGGGCAGCAAACGGAACTGAGGTACCAGAGTGAGAACGAGT |
| | GAGGAGGGCAGCAAACGGAATCCAGTTTCTGGGTTACGACCACCG |
| | GAGGAGGGCAGCAAACGGAACAGGACTCGTCGTAGTGAGCATTTT |
| | GAGGAGGGCAGCAAACGGAATGGTCCACATCGATGCGGCATAACC |
| | GAGGAGGGCAGCAAACGGAAGTCACCAGAAGCCCAAACAAAGATG |
| | GAGGAGGGCAGCAAACGGAACCCTTAACAATAGCTCTCATTGTAG |
| | GAGGAGGGCAGCAAACGGAACATCGGTTGGTCCCCAGCTGGCAGA |
| | GAGGAGGGCAGCAAACGGAACATGGAGTTAGCCATGCTGTCTGGA |
| | GAGGAGGGCAGCAAACGGAACTGGCTGCAGCAATTTCTCCAGCAC |
| | GAGGAGGGCAGCAAACGGAACGGTGTATCGTGGGTATGGGAATGG |
| | GAGGAGGGCAGCAAACGGAAAGCGTTAAAGTTATTCTTCAGATCT |
| | GAGGAGGGCAGCAAACGGAACCTTCTCCACATTAAGATCTAGACG |
| | GAGGAGGGCAGCAAACGGAACAGATACCATTGATATTGGTAGAGA |
| | GAGGAGGGCAGCAAACGGAATCAAATTGTGTTTGAAGACGTTCGG |
| | GAGGAGGGCAGCAAACGGAATGTATCCAGTGAGTTGCTCAACGTG |
| | GAGGAGGGCAGCAAACGGAAATGATGTCCAACTGATCGACGAGTT |
| | GAGGAGGGCAGCAAACGGAATGATATTCGGTATCACTTCCCAACA |
| | GAGGAGGGCAGCAAACGGAATAGCAATCCGATGGGCATTCTCGAT |
| | GAGGAGGGCAGCAAACGGAAATTAGTAAAAACCTCTACTGCCTCA |
| | GAGGAGGGCAGCAAACGGAAAGCCAGAATGGATTCCGGCATTGCA |
| | TTAATGAGTCATACGCTTATGAGCCTAGAAGAGTCTTCCTTTACG |
| | TGTCCCATGAGTGGCTCGATGCCCGTAGAAGAGTCTTCCTTTACG |
| | TAAGAGTGAACTTCTTCAGTGTTCCTAGAAGAGTCTTCCTTTACG |
| | AACCAATCTCCACTTTCCTCTTGGATAGAAGAGTCTTCCTTTACG |
| | GGCCAGTTAGTGAAACCATCCTTTGTAGAAGAGTCTTCCTTTACG |
| | TCATTGTGCGTGTTCCCATGGGTGATAGAAGAGTCTTCCTTTACG |
| | AGTTGAGTTGAAAGAGATAACAGCTTAGAAGAGTCTTCCTTTACG |
| | GCTGAGTCGGCACAGCCATCGGTTTTAGAAGAGTCTTCCTTTACG |
| | GAATCTCATGAGGAACATCGATCGTTAGAAGAGTCTTCCTTTACG |
| | AGTTTTCCAGACCATAGCCAGCATATAGAAGAGTCTTCCTTTACG |
| | CCGAACAAATGGTTGTATTCCAAACTAGAAGAGTCTTCCTTTACG |
| | TACAGTTGCCAGTAGACTTCTTGTGTAGAAGAGTCTTCCTTTACG |
| | AAGATCACGACATCTGCCATCGAACTAGAAGAGTCTTCCTTTACG |
| | AAATGCTGGAGATCTCTCCAGGTCATAGAAGAGTCTTCCTTTACG |
| | AAACTCCGGCTGCTTCCGGAGCAGCTAGAAGAGTCTTCCTTTACG |
| | ACGTCCGTATAAGTCAGTGGTAGCTTAGAAGAGTCTTCCTTTACG |
| | GAAAATGTGGAAGCCAGAGTTGAAGTAGAAGAGTCTTCCTTTACG |
| | CATTGTTGATGGCTGAATTGATGGATAGAAGAGTCTTCCTTTACG |
| | ACAATTGCAATCATCATCCTCTCCTTAGAAGAGTCTTCCTTTACG |
| | CCCAAGCCTTCGCGTCCCTCATTGATAGAAGAGTCTTCCTTTACG |
| | TACGAGGTCCATCAACAGTCTTGCCTAGAAGAGTCTTCCTTTACG |
| | AATATGGATTTTGCTTGGCTCATGTTAGAAGAGTCTTCCTTTACG |
| | ACTCCAACTCCACAGACTCCGTTGTTAGAAGAGTCTTCCTTTACG |
| | TAGTTCCATGAGAGTTGAACCAGTCTAGAAGAGTCTTCCTTTACG |
| | ATTGGATGAGAAGTCATAACTGGCTTAGAAGAGTCTTCCTTTACG |
| | TTTTTCCGGTAAATCCGAGAGCCCATAGAAGAGTCTTCCTTTACG |
| | TTTTCCTCCAGCTTGGCCAGTGTTCTAGAAGAGTCTTCCTTTACG |
| | TCAGAGGGTGATTGAACGGATGAGATAGAAGAGTCTTCCTTTACG |
| | GAGGTCTGTATCCTCGTTTTGTTCGTAGAAGAGTCTTCCTTTACG |
| | GACCTCATTATCCTTGTGGAGCTTGTAGAAGAGTCTTCCTTTACG |
| | GCATGTGATAAAGCTGGTTGAATAATAGAAGAGTCTTCCTTTACG |
| | GTCCTCTATTTATGAATCCGTGTCGTAGAAGAGTCTTCCTTTACG |
| | GGGTTCCTTTACATGAACATGAAAATAGAAGAGTCTTCCTTTACG |
| | AGAGACAAGATTACCAGAAAGAGGCTAGAAGAGTCTTCCTTTACG |
| **B3_*Ttu_ast-1*** | GTCCCTGCCTCTATATCTTTATTAGAGCCGTAAAGTGGAAAGTTT |
| | GTCCCTGCCTCTATATCTTTGAAAAACTACCATATCCCATACTTG |
| | GTCCCTGCCTCTATATCTTTAGCCGGGTTGGATGAGAGAACGATA |
| | GTCCCTGCCTCTATATCTTTAGGTAATCGGTTGACAGCCAGATAA |
| | GTCCCTGCCTCTATATCTTTTGAATAGCAGCGGCAATTCCTTGAA |
| | GTCCCTGCCTCTATATCTTTGAACCTTGGCCATGATATTCTTGTC |
| | GTCCCTGCCTCTATATCTTTATCGTAATTCATGTTCGGTTTCGAT |
| | GTCCCTGCCTCTATATCTTTTCCGGATCAACAAGTTTGAATTCGC |
| | GTCCCTGCCTCTATATCTTTGATGACGAGCATCTGACAGCAATTC |
| | GTCCCTGCCTCTATATCTTTACTGCCTGAGTGAGCCAATTGTCTG |
| | GTCCCTGCCTCTATATCTTTTGAGTACTGAAAGAGCTTGGATTGC |
| | GTCCCTGCCTCTATATCTTTTGAAGCATGGCATGTTGAATTGAGG |
| | GTCCCTGCCTCTATATCTTTAGCTCCAGCTCCAGCTGCTGCAGCA |
| | GTCCCTGCCTCTATATCTTTTGCATCGAGAGATTCATACCCGAGA |
| | GTCCCTGCCTCTATATCTTTGAGCCATCGTTCCACTGTTTGATTC |
| | GTCCCTGCCTCTATATCTTTTCCATTGGGCACACCGTTCTGTAGA |
| | GTCCCTGCCTCTATATCTTTACAGCAACGGCTGCGCTAGCAACTG |
| | GTCCCTGCCTCTATATCTTTCGGATCCGTTTCCTTCCCTTTTGGG |
| | GTCCCTGCCTCTATATCTTTACCATATTCAATAACAGCCGACCAG |
| | TTATGCACATTTGGAGTAATTGCTCTTCCACTCAACTTTAACCCG |
| | GTAGGTAATGATGTATTGTTGGCTGTTCCACTCAACTTTAACCCG |
| | GGGTGAAGAGAGCTGTACCCGGTTGTTCCACTCAACTTTAACCCG |
| | GCTCCATGGACCCTGCATGAATTCTTTCCACTCAACTTTAACCCG |
| | GCCGACGTTGAGCTGGGTGTTTGAGTTCCACTCAACTTTAACCCG |
| | CGAATTTGTAAGCATATCGTTTTCCTTCCACTCAACTTTAACCCG |
| | ATAATATCTTAGAGCACGAGACATCTTCCACTCAACTTTAACCCG |
| | CGTTCTCCCCATTTTCTGGCAACCTTTCCACTCAACTTTAACCCG |
| | GAGTTCCTTCCCAAGTGATAACTTCTTCCACTCAACTTTAACCCG |
| | CAGGAATTGCCATAATTGAGTCTGATTCCACTCAACTTTAACCCG |
| | GTAGTACAAAGAGTCTGGTAAGGATTTCCACTCAACTTTAACCCG |
| | TTGGAATAACATGCTTCAAGTTATATTCCACTCAACTTTAACCCG |
| | TTGAAGAGTCTCATCTTTAACTTGATTCCACTCAACTTTAACCCG |
| | GCAGAAACAGCCGAACTGTTGAAGGTTCCACTCAACTTTAACCCG |
| | CATCAGAACCTGAGTTTTTGGTTGTTTCCACTCAACTTTAACCCG |
| | AAAACTGGAAAGACCTCGAGGAAGATTCCACTCAACTTTAACCCG |
| | GGAAAGGAAACTGCTTCAGCAGCTTTTCCACTCAACTTTAACCCG |
| | CGGCGTTTTCCGTCTTGACGTCCATTTCCACTCAACTTTAACCCG |
| | AAATCGCTGCTCTATGTTAACAGACTTCCACTCAACTTTAACCCG |

**Supplemental Table 2 continued. Gene and probe sequences used in study.**

| | |
|---|---|
| B4_*Ttu_sem-4* | CCTCAACCTACCTCCAACAAACTAGCTGCTGAGGATGGCGTCGTA |
| | CCTCAACCTACCTCCAACAACCGTCCTCCATAAAAGCATCATCAT |
| | CCTCAACCTACCTCCAACAAATAGGTGACACGACGACGTCTACAG |
| | CCTCAACCTACCTCCAACAATTTCAGATTCCCTCGGGTGGTGAAG |
| | CCTCAACCTACCTCCAACAAGCTTGAAGCAGACACCACATTGATG |
| | CCTCAACCTACCTCCAACAAGTCAGCCGAGGAGATGAGGGAGCTA |
| | CCTCAACCTACCTCCAACAATGTCCTGGCAGTCGAGGTGTTCAGA |
| | CCTCAACCTACCTCCAACAACTTCACTACTGGCTGATGGAGAAAT |
| | CCTCAACCTACCTCCAACAACTCGTAGTTGGCTTCACCCGAATCT |
| | CCTCAACCTACCTCCAACAAACCGGATCCAGAAAAGGACTGAGGA |
| | CCTCAACCTACCTCCAACAATTAGCGCTTGTCGATGTTCAGCCAT |
| | CCTCAACCTACCTCCAACAATCTGTAAGAAAAATTACAAATAGTT |
| | CCTCAACCTACCTCCAACAATGGGACATTGCTGTTGTGGCTGCAG |
| | CCTCAACCTACCTCCAACAATGGCATCTGAGGCATCTGTGTGTGT |
| | CCTCAACCTACCTCCAACAACGACAAAACTCGTCGACAGAGGATG |
| | CCTCAACCTACCTCCAACAATTGTCCCCAACTAAAGCACGGATCT |
| | CCTCAACCTACCTCCAACAATCAAAGCTTCCCAGTCGTCCTCATT |
| | CCTCAACCTACCTCCAACAATGAAGGAGGAAACATGTCGACTCTC |
| | CCTCAACCTACCTCCAACAAGGAGCAGACGGATGTCCGAATGGGT |
| | CCTCAACCTACCTCCAACAATGGGCGTGACAGGAGCACCGCTGGC |
| | CCTCAACCTACCTCCAACAAACCAGCAGCTGCAGCTACTGAAGCC |
| | CCTCAACCTACCTCCAACAATTCAGTGAGCCATTTGTGAGACCCA |
| | CCTCAACCTACCTCCAACAAGTTGAAGCTGCTGAAGTTGCTGAAG |
| | CCTCAACCTACCTCCAACAAGTGTAGATTATGATTGTTGGTGGAT |
| | CCTCAACCTACCTCCAACAATGTTGAGCAAAAAGCTGGTTGAGCA |
| | CCTCAACCTACCTCCAACAACAGTACCATCCTCATCAGCCATCGA |
| | CCTCAACCTACCTCCAACAAAACAATGGAAAAGCTTGGTCCAGCT |
| | CCTCAACCTACCTCCAACAATCAGGACTGTTGACAGTCGATCGTT |
| | CCTCAACCTACCTCCAACAAGTTCACACTCATGTTCAATAATCTG |
| | CCTCAACCTACCTCCAACAAGTCGCATTGTAATAGCTTTTCTGTT |
| | CCTCAACCTACCTCCAACAAGCACCTTGTGAGCTGAAGCACTGTC |
| | CCTCAACCTACCTCCAACAACAGCAGTCGAATCATCGACCATCAT |
| | CCTCAACCTACCTCCAACAATAAAGATTGATGTTGCTGTCGTTGA |
| | CCTCAACCTACCTCCAACAATGCTGAGGTGTAGCTATGGATGCTA |
| | CCTCAACCTACCTCCAACAACATCTAGCATTTTGTCCATCAAGGC |
| | CCTCAACCTACCTCCAACAAATCAATCGGCGACGACGACTCGTCC |
| | TTATTCGGCAGCAGTAGTAGGTTGCATTCTCACCATATTCGCTTC |
| | CACAAACTTTTTGACAAACAGAGCAATTCTCACCATATTCGCTTC |
| | AAGCTGAGAGATGCGGAAGAGCCTTATTCTCACCATATTCGCTTC |
| | TTGCCACATATGAGTACCCATATGAATTCTCACCATATTCGCTTC |
| | TCTGGAGAGCTGAGCTTGAGCTGAAATTCTCACCATATTCGCTTC |
| | TGCATTGCTTCTAATGGATTATCTGATTCTCACCATATTCGCTTC |
| | GTCGGAAGAGGAGGATTTGCTGGTGATTCTCACCATATTCGCTTC |
| | AAGAACCGTCTGTTTGATCATGGGAATTCTCACCATATTCGCTTC |
| | GGGGGAGGACTTCCAGTAGTGGATAATTCTCACCATATTCGCTTC |
| | AGCTCCTACTCCAGCCCCGGTCTGCATTCTCACCATATTCGCTTC |
| | GCGGCATTCCACTGGGTATCCGTCCATTCTCACCATATTCGCTTC |
| | CGCTTGAAGTTGCGAAGGTGTCATAATTCTCACCATATTCGCTTC |
| | TCAAAATGCTGAACCTCTTCTGGCAATTCTCACCATATTCGCTTC |
| | AGCCATTGACAAATGATTTCCCAACATTCTCACCATATTCGCTTC |
| | GTAATGCATCTGAAGTGCTGACTTGATTCTCACCATATTCGCTTC |
| | TGATTGGGATCTGTCGTCGGCATAGATTCTCACCATATTCGCTTC |
| | CGGCTTCATCCGTTGTCGAAATCTCATTCTCACCATATTCGCTTC |
| | GATTTCCGGTCGAACTCCACCAGCAATTCTCACCATATTCGCTTC |
| | CCATCAAATGGTCTCATGAAAAAGGATTCTCACCATATTCGCTTC |
| | GCGATAATGGAAACATGGCTGCCGAATTCTCACCATATTCGCTTC |
| | GGTTGGCGTCTGATTGTCGCCAGAAATTCTCACCATATTCGCTTC |
| | TGAGGACCGATGGAGTAATTGTCAGATTCTCACCATATTCGCTTC |
| | GATGTTGTTGTTGTTGATTTTGAAGATTCTCACCATATTCGCTTC |
| | CTGCGCATTTTGAATATGTTGCTGAATTCTCACCATATTCGCTTC |
| | TTGGATGACGAATTGATGGAGACATATTCTCACCATATTCGCTTC |
| | TGGCTGAGGAAGGTAAGAGCATGGCATTCTCACCATATTCGCTTC |
| | GCTCAAGGCAAGATCGTCTTCAGCTATTCTCACCATATTCGCTTC |
| | GCTGGAACAGATCCACGATCAGCAGATTCTCACCATATTCGCTTC |
| | CACCAGCTGATCGTGAACGCATCTGATTCTCACCATATTCGCTTC |
| | ACGATCGCTAAAACGTTGTCCACATATTCTCACCATATTCGCTTC |
| | TGTTCTTCCATCAAATGCCACAGATATTCTCACCATATTCGCTTC |
| | TTTTTGGTGTTGCCGGAGGCGTTACATTCTCACCATATTCGCTTC |
| | CACCTCATGCTCCTGCTCGGTTGACATTCTCACCATATTCGCTTC |
| | GCTAATTGAGCGCACAATGTTAAAAATTCTCACCATATTCGCTTC |
| | ATTGGAGCAACAAGGATTTTCCGTTATTCTCACCATATTCGCTTC |
| | GGCCGAGCTTGCCGTCGTCGTCGACATTCTCACCATATTCGCTTC |
| B1_*Ttu_nlp-3* | GAGGAGGGCAGCAAACGGAAAGGATAGAATAGGATTTGCCTTCCG |
| | GAGGAGGGCAGCAAACGGAAGGTTTGCCATTCTCTGAGAAGGTAC |
| | GAGGAGGGCAGCAAACGGAAAGACGGCTCCTAGGATGAGAAGAGC |
| | GAGGAGGGCAGCAAACGGAATGGCAAAATATTTGGATCTCGTGCT |
| | GAGGAGGGCAGCAAACGGAACTGAAATAAGAGCCTTAGGAAGCCA |
| | TTTTTCCATGTTAAAAGTGGCTTCATAGAAGAGTCTTCCTTTACG |
| | TTTCCAGAGATCATATATCCCATAATAGAAGAGTCTTCCTTTACG |
| | TACGCTCTTCAGCTTGAACAATGCATAGAAGAGTCTTCCTTTACG |
| | GAAAACTAGCTTCATATTGGCCGGCTAGAAGAGTCTTCCTTTACG |
| | TTTTTCGTCCACGGCCCCATCTTGATAGAAGAGTCTTCCTTTACG |

**Supplemental Table 2 continued. Gene and probe sequences used in study.**

### Supplemental Video Legends

**Supplemental Video 1. *Tokorhabditis tufae*, unlike most nematodes, gives live birth to progeny.** Real-time recording of a day-2 adult.

**Supplemental Video 2. Vulval contractions and twitching precede *C. elegans* egg laying but not *T. tufae* live birth.** Real-time recording of *C. elegans* egg laying and *T. tufae* live birth. Animals are day-2 adults.

**Supplemental Video 3. *T. tufae* larva during live birth.** Representative high-speed recording of a day-2 adult showing larval movement at birth.

**Supplemental Video 4. Dissected *T. tufae* larvae recapitulate the transition from quiescence to activity.** Representative sped-up recording of four *T. tufae* dissected larvae from a day-2 adult. Larvae transition from quiescence to activity at 00:27, 02:38, 05:16, and 06:28 (hour:min).

**Supplemental Video 5. *T. tufae* Unc larvae and adults demonstrate uncoordinated movement.** Real-time recording of live birth from wild type and Unc day-2 adults.

**Supplemental Video 6. Live birth is observed after adults are killed by SDS.** Real-time recording of *T. tufae* live birth from viable and SDS-killed day-2 adults.
