## Supplementary figures and images for "Live birth protects the developing progeny of the extremophile nematode *Tokorhabditis tufae*"

### Supplemental Video 1

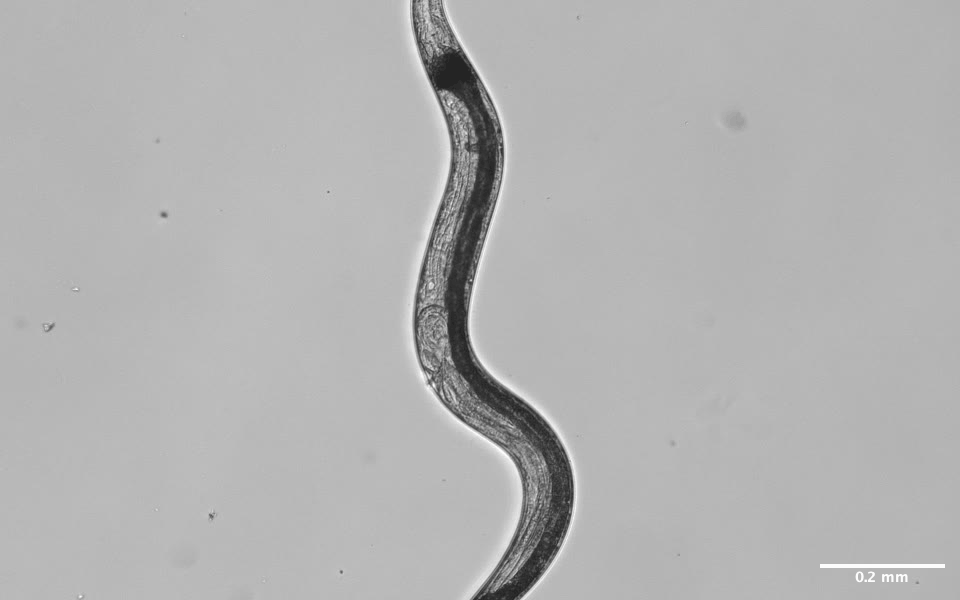
